# Unveiling the Biochemical Mechanisms of Aging and the Implications of Oxidative Stress on Cellular Senescence through Multi-Omics Analysis of Fibroblasts

**DOI:** 10.1101/2025.05.17.654671

**Authors:** Rajarshi Mandal, Ning Xie, Gil Alterovitz

## Abstract

This research investigates the complex biochemical mechanisms underlying aging by analyzing primary human fibroblasts using a longitudinal multi-omics dataset. This dataset includes cytology, DNA methylation and epigenetic clocks, bioenergetics, mitochondrial DNA sequencing, RNA sequencing, and cytokine profiling. Key findings indicate that mitochondrial efficiency declines with age, while glycolysis becomes more prevalent to compensate for energy demands. Epigenetic clocks, such as Hannum and PhenoAge, showed strong correlations with biological age (ρ > 0.650, p < 1e-6), validating the experimental setup and confirming that the cultured fibroblasts were aging appropriately. Fibroblasts with SURF1 mutations exhibited accelerated aging, marked by bioenergetic deficits, increased cell volume, and reduced proliferative capacity, underscoring the pivotal role of mitochondrial dysfunction in cellular senescence. Novel insights were gained from analyzing cytokines like IL18 and PCSK9, some of which were linked to age-related diseases such as Alzheimer’s and cardiovascular disorders.

Experimental treatments revealed distinct effects on cellular aging. Dexamethasone reduced inflammation but also increased DNA methylation, induced metabolic inefficiencies, and shortened cellular lifespan. Oligomycin heightened oxidative stress and RNA degradation, emphasizing how such treatments contribute to cellular stress and metabolic imbalance while shedding light on aging mechanisms. By uncovering connections between mitochondrial dysfunction, epigenetic biomarkers, and immune dysregulation, this study identifies potential therapeutic targets for age-related diseases. Future research could validate the most promising biomarkers across diverse cell types and experimental treatments to build a more comprehensive understanding of aging.

## Introduction

Biogerontology is a field that focuses on understanding the underlying mechanisms of aging to find potential interventions to promote longevity. Currently, there is no scientific consensus on a particular aging theory, but they have been classified into two major categories: programmed aging theories and damage theories [1]. Programmed aging theories propose that aging is genetically regulated and results in evolutionary advantages such as promoting genetic adaptation and removing post-reproductive age individuals to combat overpopulation [2]. In contrast, most evolutionary biologists believe that the lack of natural selection in post-reproductive organisms leads to reduced maintenance.

One promising explanation is the free radical theory of aging, which suggests that reactive oxygen species (ROS) damage biomolecules such as DNA, protein, and lipids. This causes an accumulation of structural and functional problems inside cells [3]. ROS are highly reactive chemicals created in metabolic reactions. Common ROS include hydroperoxide, superoxide, hydroxyl radical, and singlet oxygen and these can cause oxidative stress.

Researchers at MRC Laboratory of Medical Sciences extended the lifespan of *Drosophila melanogaster* by amplifying expression of superoxide dismutase (SOD) and catalase, which are antioxidant enzymes [4]. Mitochondria are major producers of ROS in mammalian cells, which leads to mitochondria DNA (mtDNA) being highly vulnerable to oxidative stress. After realizing that the lack of mitochondrial homeostasis is a shared hallmark among numerous age-associated ailments, the free radical theory has been updated to focus on the clonal amplification of pre-existing mtDNA mutations from reduced mitochondrial polymerase γ fidelity [5].

Epigenetics is the study of heritable or stable changes that occur without changes to the DNA sequence; some well-known examples include DNA methylation, histone modification, and non-coding RNA regulation [6]. DNA methylation is the biological process where methyl groups are added to a DNA molecule, which has been shown to cause genomic imprinting, X-chromosome inactivation, and repression of transposable elements [7, 8]. Recently, it has been found to strongly correlate with biological age through epigenetic clocks, which are biochemical tests that quantify biological age from CpG site methylation data. The most renowned clocks include Horvath, Hannum, PhenoAge, and GrimAge; each has their own unique architecture and differentiating factor [9].

Researchers at Columbia University created a longitudinal dataset with high temporal resolution from cultured primary human fibroblasts measured across their replicative lifespans [10], which we thoroughly analyzed in this study. A fibroblast is a cell that helps connective tissue form through secreting collagen proteins that maintain structure and provide support.

These were isolated from skin of six healthy donors and dermal punch biopsies of three individuals with a lethal variant of SURF1 mutation (IRB #AAAB0483). Surfeit locus protein 1 (SURF1) is part of the mitochondrial translation regulation assembly intermediate of cytochrome c oxidase (COX) complex [11]. Mutations in its encoding gene have been associated with Leigh syndrome, a neurological condition that affects the central nervous system, and Charcot–Marie–Tooth disease [12].

The dataset from the Columbia University researchers encompasses various molecular and cellular data types, including cytology, DNA methylation and epigenetic clocks, bioenergetics, mitochondrial DNA sequencing, RNA sequencing, and cytokine profiling. Some of the variables include total days grown, days after treatment, clinical condition, and treatment types. The “Clinical Condition” refers to whether the fibroblast has the SURF1 mutation while the “Treatments” refers to the name of the experimental treatment. Some treatments in the dataset include Dexamethasone, Oligomycin, Mitochondrial Nutrient Uptake Inhibitors + Dexamethasone, Mitochondrial Nutrient Uptake Inhibitors, 5-azacytidine, Contact Inhibition, 5-azacytidine + Mitochondrial Nutrient Uptake Inhibitors, Galactose, 2-Deoxy-D-glucose, and betahydroxybutyrate. Due to a low amount of data in some categories for most treatments, we analyzed the effects of only Dexamethasone and Oligomycin in this paper. Dexamethasone, a corticosteroid fluorinated at position 9, is most frequently used for treating inflammation, arthritis, severe allergies, and asthma because it is a glucocorticoid receptor agonist [13, 14]. It is similar to a natural hormone produced by human adrenal glands and was even found to reduce mortality in patients with severe COVID-19 [15]. Oligomycin, a macrolide produced by bacteria in the genus *Streptomyces*, inhibits ATP synthase (Complex V) by blocking the F_O_ subunit of the proton channel, halting oxidative phosphorylation. However, researchers are looking into its use as an antibiotic [16].

The control group in the dataset was the cells without the SURF1 mutation cultured without any experimental medium. The experimental groups included cells with the SURF1 mutation and healthy cells cultured with various treatments like Dexamethasone and Oligomycin. We tested a plethora of hypotheses in this study. For example, we hypothesized that DNA methylation and mitochondrial DNA sequencing biomarkers’ correlation with biological age would support the free radical theory of aging. We also hypothesized that fibroblasts with the SURF1 mitochondrial mutation would exhibit exacerbated oxidative stress and impaired bioenergetics, validating mitochondrial dysfunction as a critical driver of senescence. Finally, we hypothesized that experimental treatments such as Dexamethasone and Oligomycin would emulate various conditions so that we could better understand how certain biological pathways are impacted as we age.

## Methods

### Data Preprocessing

The original dataset is available at the <u>Columbia Picard Shiny App</u>. First, the dataset was downloaded from https://columbia-picard.shinyapps.io/shinyapp-Lifespan_Study/ in the “Download Data” subpage with the checkbox selected for “Click to download the full sample-set.” After it was uploaded to Kaggle at https://www.kaggle.com/datasets/rm1000/multi-omics-aging-with-mitochondrial-perturbations, a Jupyter Notebook running Python 3.12 was created.

Next, we imported the essential libraries for data preprocessing including math, NumPy, pandas, matplotlib, seaborn, SciPy, and warnings. The csv file containing the original data was read using pandas’ read_csv function and the column names were manually renamed for clarity and conciseness.

### Data Visualization

Next, we wanted to visualize some biomarker measurement distributions from each of the six categories mentioned earlier (samples in Fig. 1). To achieve this, we chose a column for each category and plotted them using seaborn’s histplot function with 16 bins. In other words, the x-axis contains ranges of values for the chosen columns while the y-axis displays the percentage of datapoints that belong to the corresponding range of values.

**Figure 1:**
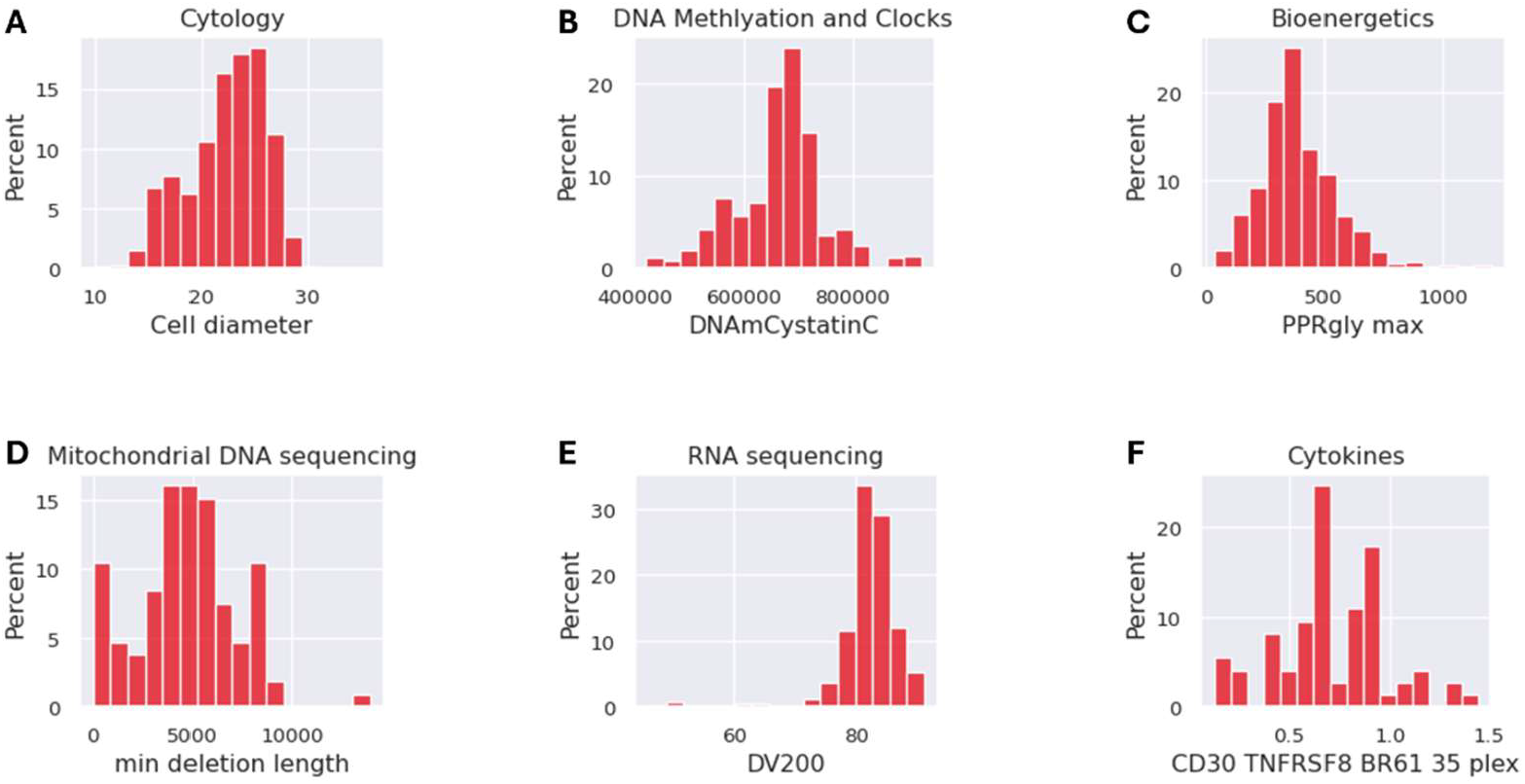
Visualizing dependent variables (biomarkers) with histograms. **A.** The histogram shows the distribution of cell diameter values. **B.** The histogram displays the amount of DNA methylation on the encoding gene of Cystatin C. **C.** The histogram shows the maximum measured PPR glycolysis. **D.** The histogram shows the minimum deletion length in mitochondrial DNA. **E.** The histogram displays DV200, a metric used to quantify the quality of RNA by measuring the percentage of fragments over 200 nucleotides. **F.** The histogram shows the amount of TNF receptor superfamily member 8, a cell membrane protein, across all measurements.

**Figure 2:**
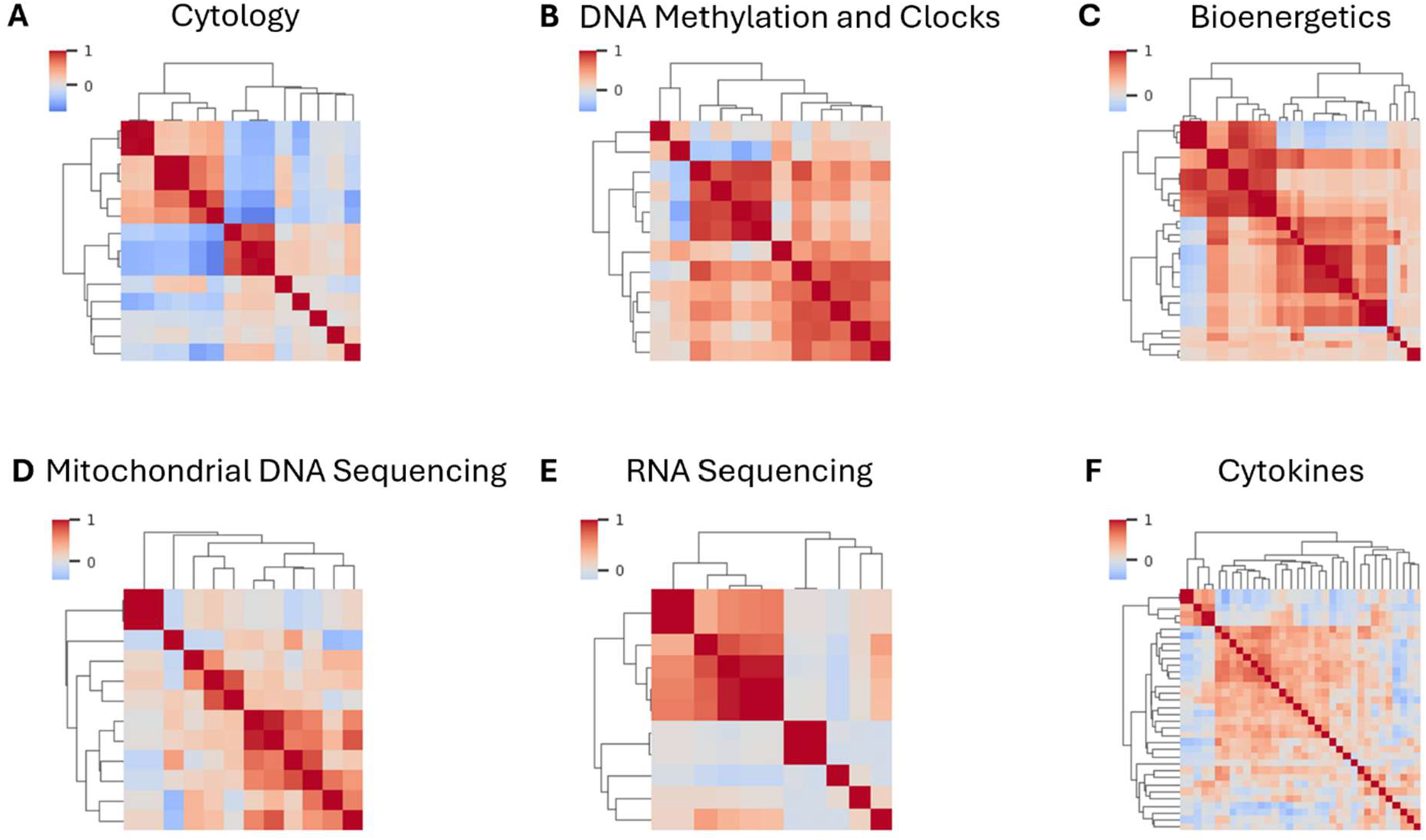
Revealing correlation between dependent variables with clustermaps. **A.** There is a moderate amount of positive and negative correlations for cytology columns. **B.** There is a significant positive correlation between DNA methylation and clocks columns. **C.** There is a significant positive correlation between bioenergetics columns. **D.** There is little correlation between mitochondrial DNA sequencing columns. **E.** There is a strong positive correlation between some RNA sequencing columns. **F.** There is a weak positive correlation between some cytokines.

**Figure 3:**
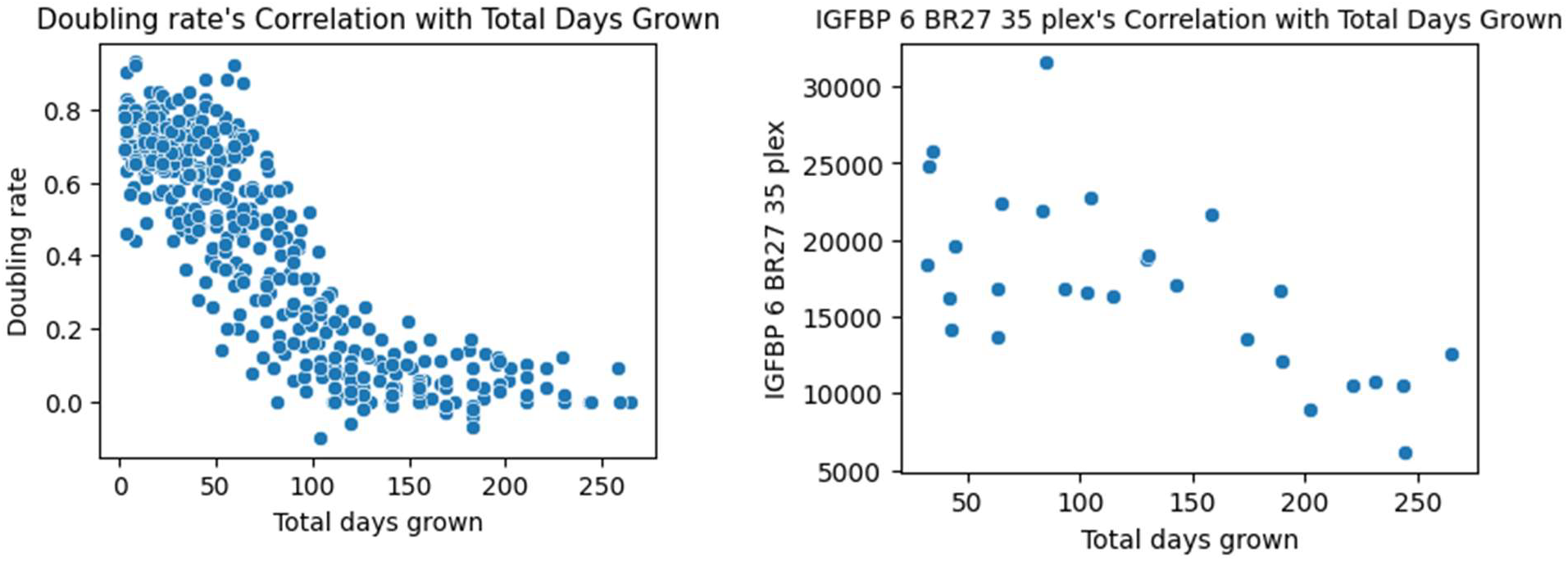
Visualizing biomarker measurements over age. The scatterplot on the left plots doubling rate against total days grown and there is a clear negative relationship. The scatterplot on the right graphs insulin-like growth factor-binding protein 6 and total days grown; there is a weak negative relationship.

**Figure 4:**
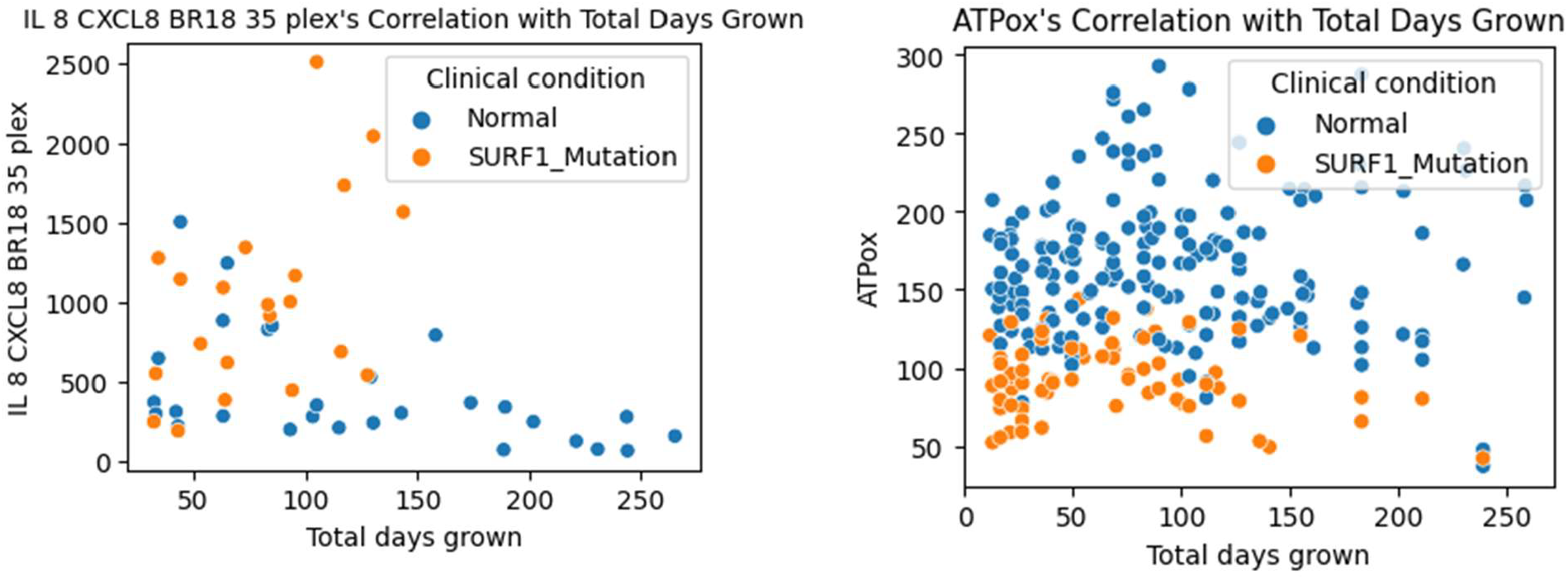
Impact of SURF1 mutation on biomarkers. The graph on the left indicates that the SURF1 mutation positively impacts the amount of interleukin 8. The scatterplot on the right implies that the SURF1 mutation negatively impacts the amount of ATP oxidation in the cultured fibroblasts.

**Figure 5:**
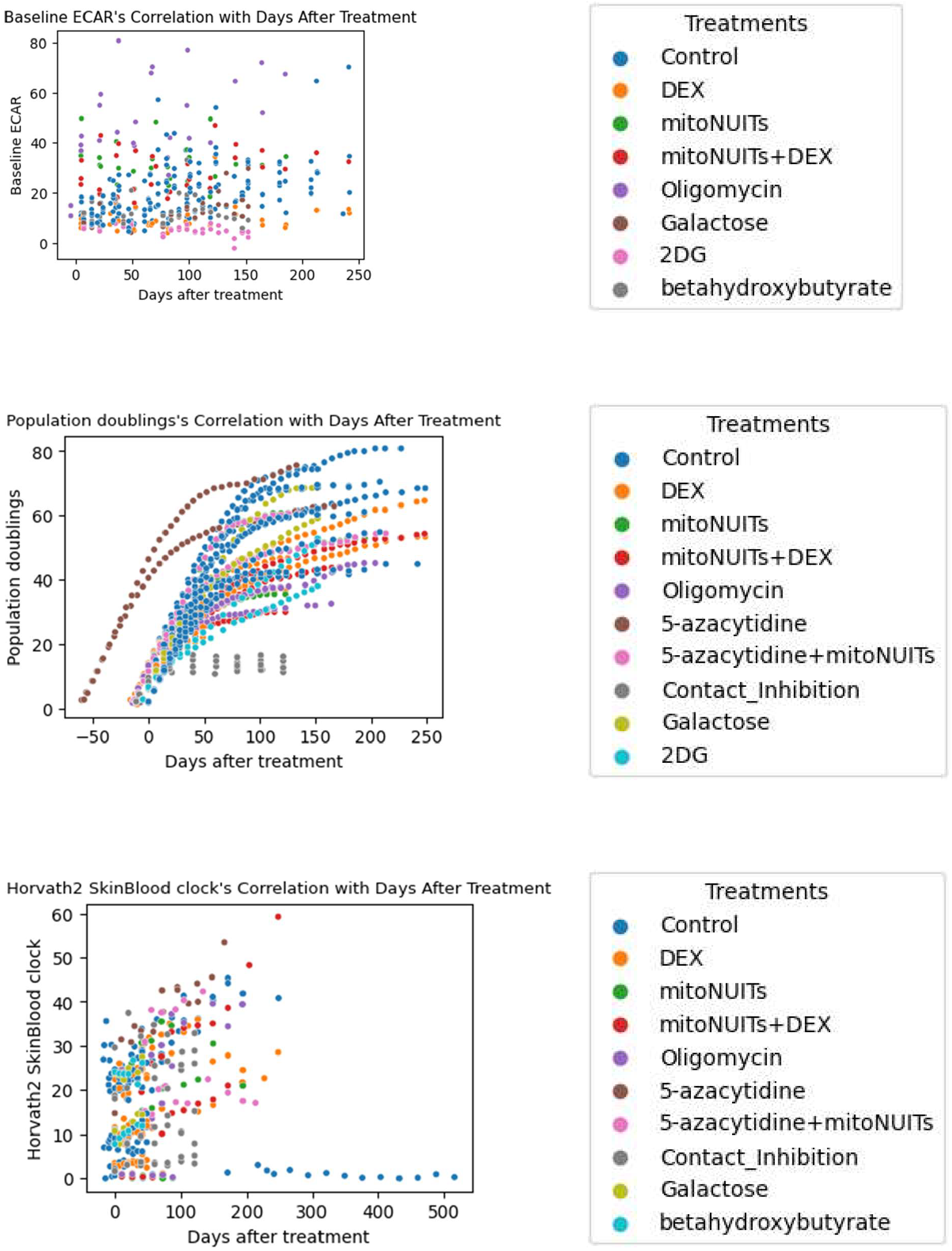
Impact of experimental treatments on biomarkers. The top scatterplot shows the effect of 7 treatments on the baseline extracellular acidification rate. The middle scatterplot shows the effect of 9 treatments on population doublings. The bottom scatterplot shows the effect of 9 treatments on the Horvath2 skin and blood epigenetic clock.

After that, we visualized the correlations between the columns in each of the categories. Clustermaps were perfect for this because they plotted matrix data as a hierarchically clustered heatmap. The dark red indicates a strong positive correlation, the grey indicates no correlation, and the dark blue indicates a strong negative correlation. The column names are not included because there were too many of them to display.

### Statistical Tests

First, we identified treatments correlated with the number of days grown after treatment by running student’s t-test for paired samples on the experimental treatments and the number of days grown after treatment. For each of the biomarker categories, the following statistical tests were run.

To figure out which biomarkers were predictors of total days grown, we ran Spearman’s rank correlation between the biomarkers and total days. This is because there are two quantitative variables that are nonparametric. The data corresponding to this statistical test was visualized using scatterplots with the total days grown on the x-axis and the biomarker on the y-axis.

Next, we ran student’s t-test for paired samples between the biomarkers and whether or not the fibroblast has the SURF1 mutation to determine which biomarkers were impacted by the SURF1 mutation. This is because there is one quantitative and one qualitative variable with two paired groups that are parametric. The data corresponding to this statistical test was visualized using scatterplots with the total days grown on the x-axis, the biomarker on the y-axis, and whether the fibroblast had the SURF1 mutation as the color of the point.

To figure out which biomarkers were impacted by each treatment, we ran student’s t-test for paired samples between the biomarkers and whether or not the specific treatment had been used. This is because there is one quantitative and one qualitative variable with two paired groups that are parametric. We also created scatterplots with the days after treatment on the x-axis, the biomarker on the y-axis, and the treatment as the color of the point. Three example scatterplots with their legends are shown below.

## Results

In each of the data tables, the first column has the category, the second column has the biomarker measurement, the third column has the p-value, and the fourth column the result of the statistical test run using the Python library SciPy rounded to 3 decimal points if the p-value is less than 0.05. This means that the particular result was statistically significant, and the null hypothesis can be rejected with 95% confidence. The first data table contains the result of Spearman’s rank correlation between the dependent variables (biomarkers) and total days grown.

**Table 1:** Quantifying correlation between biomarkers and biological age.

| Category | Biomarker | p-value | $\rho$ |
| --- | --- | --- | --- |
| Cytology | Population doublings | < 0.001 | 0.941 |
|  | Days per passage | < 0.001 | 0.669 |
|  | Population doubling time | < 0.001 | 0.598 |
|  | Percent dead cells | < 0.001 | 0.419 |
|  | Cells plated | < 0.001 | 0.339 |
|  | Cell volume | < 0.001 | -0.329 |
|  | Cell diameter | < 0.001 | -0.329 |
|  | Cells counted | < 0.001 | -0.541 |
|  | Live cells counted | < 0.001 | -0.546 |
|  | Divisions per passage | < 0.001 | -0.709 |
|  | Doubling rate | < 0.001 | -0.846 |
| DNA<br>Methylation<br>and Clocks | Mitotic age | < 0.001 | 0.839 |
|  | DNAmLeptin | < 0.001 | 0.766 |
|  | Hannum clock | < 0.001 | 0.706 |
|  | PCPhenoAge clock | < 0.001 | 0.650 |
|  | PCHannum clock | < 0.001 | 0.618 |
|  | PhenoAge clock | < 0.001 | 0.431 |
|  | PCHorvath2 SkinBlood clock | < 0.001 | 0.373 |
|  | DNAmPACKYRS | 0.007 | 0.234 |
|  | DNAmSen clock | 0.013 | 0.216 |
|  | DNAmCystatinC | 0.016 | -0.210 |
|  | DunedinPoAm rate of aging | 0.016 | -0.210 |
|  | DunedinPACE rate of aging | 0.005 | -0.244 |
|  | GrimAge clock | 0.001 | -0.291 |
|  | DNAmB2M | < 0.001 | -0.308 |
|  | DNAmADM | < 0.001 | -0.475 |
|  | DNAmPAI1 | < 0.001 | -0.596 |
|  | PCDNAmTL | < 0.001 | -0.875 |
| Bioenergetics | PPRgly | < 0.001 | 0.513 |
|  | ATPglyc | < 0.001 | 0.505 |
|  | Baseline ECAR | < 0.001 | 0.502 |
|  | PPRtotal | < 0.001 | 0.502 |
|  | Max ECAR | < 0.001 | 0.473 |
|  | PPRtotal max | < 0.001 | 0.473 |
|  | ATPglyc max | < 0.001 | 0.470 |
|  | PPRgly max | < 0.001 | 0.462 |
|  | Resting metabolic rate watts per kilogram | < 0.001 | 0.426 |
|  | ATPtotal | < 0.001 | 0.426 |
|  | Resting metabolic rate picoWatts per cell | < 0.001 | 0.426 |
|  | Max metabolic rate watts per kilogram | < 0.001 | 0.349 |
|  | Max metabolic rate picoWatts per cell | < 0.001 | 0.349 |
|  | ATPtotal max | < 0.001 | 0.349 |
|  | Glycolytic spare capacity | < 0.001 | 0.274 |
|  | Oxidative spare capacity | 0.001 | 0.251 |
|  | ATPglyc spare | 0.001 | 0.248 |
|  | ATPtotal spare | 0.002 | 0.233 |
|  | Max metabolic rate mlO2 per kilogram per min | 0.008 | 0.199 |
|  | PPRresp max | 0.008 | 0.198 |
|  | Max respiration | 0.008 | 0.198 |
|  | Coupling efficiency | 0.003 | -0.219 |
| Mitochondrial DNA sequencing | max repeat length | 0.043 | -0.297 |
| RNA sequencing | Base 30 | 0.025 | 0.227 |
|  | quality | 0.037 | 0.212 |
|  | RIN | 0.006 | -0.276 |
|  | DV200 | 0.004 | -0.287 |
| Cytokines | TGF alpha BR63 18 plex | < 0.001 | 0.674 |
|  | IL 18 IL 1F4 BR78 35 plex | < 0.001 | 0.672 |
|  | GDF 15 BR65 35 plex | 0.001 | 0.581 |
|  | CCL7 MCP 3 MARC BR37 35 plex | 0.012 | 0.470 |
|  | ALCAM CD166 BR52 18 plex | 0.036 | 0.397 |
|  | Fas TNFRSF6 BR73 35 plex | 0.047 | -0.379 |
|  | CCL23 MPIF 1 BR56 35 plex | 0.031 | -0.409 |
|  | Follistatin like.1 FSTL1 BR55 35 plex | 0.026 | -0.419 |
|  | CD30 TNFRSF8 BR61 35 plex | 0.017 | -0.448 |
|  | TFPI BR64 18 plex | 0.010 | -0.480 |
|  | X4 1BB TNFRSF9 CD137 BR13 35 plex | 0.004 | -0.527 |
|  | IL 8 CXCL8 BR18 35 plex | 0.002 | -0.562 |
|  | IGFBP 6 BR27 35 plex | < 0.001 | -0.641 |
|  | PCSK9 BR44 18 plex | < 0.001 | -0.659 |

Among all of the dependent variables (biomarkers), population doublings, mitotic age, DNAmLeptin, Hannum clock, TGF alpha BR63 18 plex, IL 18 IL 1F4 BR78 35 plex, days per passage, and PCPhenoAge clock had the strongest positive correlation with biological age.

PCDNAmTL, doubling rate, divisions per passage, PCSK9 BR44 18 plex, IGFBP 6 BR27 35 plex, DNAmPAI1, IL 8 CXCL8 BR18 35 plex, and live cells counted had the strongest negative correlation with biological age. In addition, all of these dependent variables (biomarkers) had p-values of less than 1e-6. The full results are thoroughly analyzed in the discussion section.

The next data table contains the result of student’s t-test for paired samples between the dependent variables (biomarkers) and clinical condition (normal, SURF1 mutation).

**Table 2:**
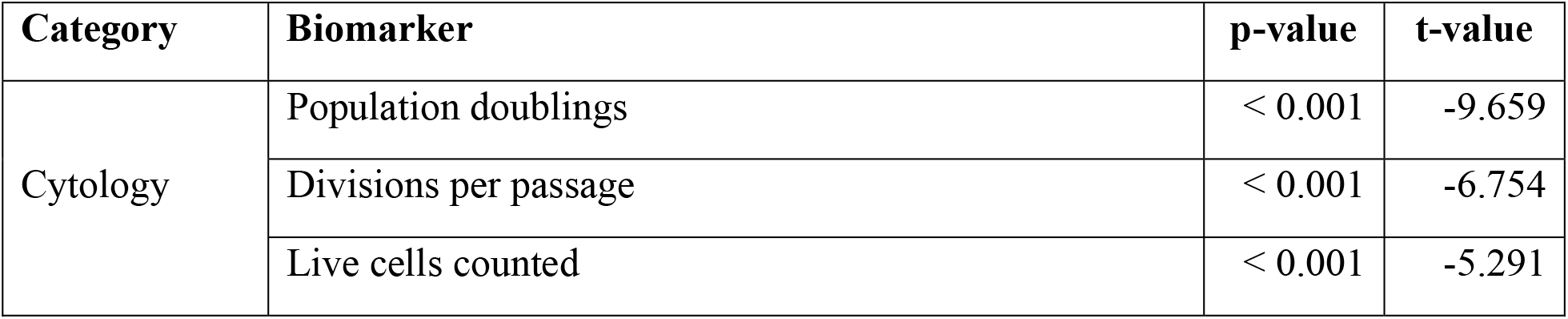
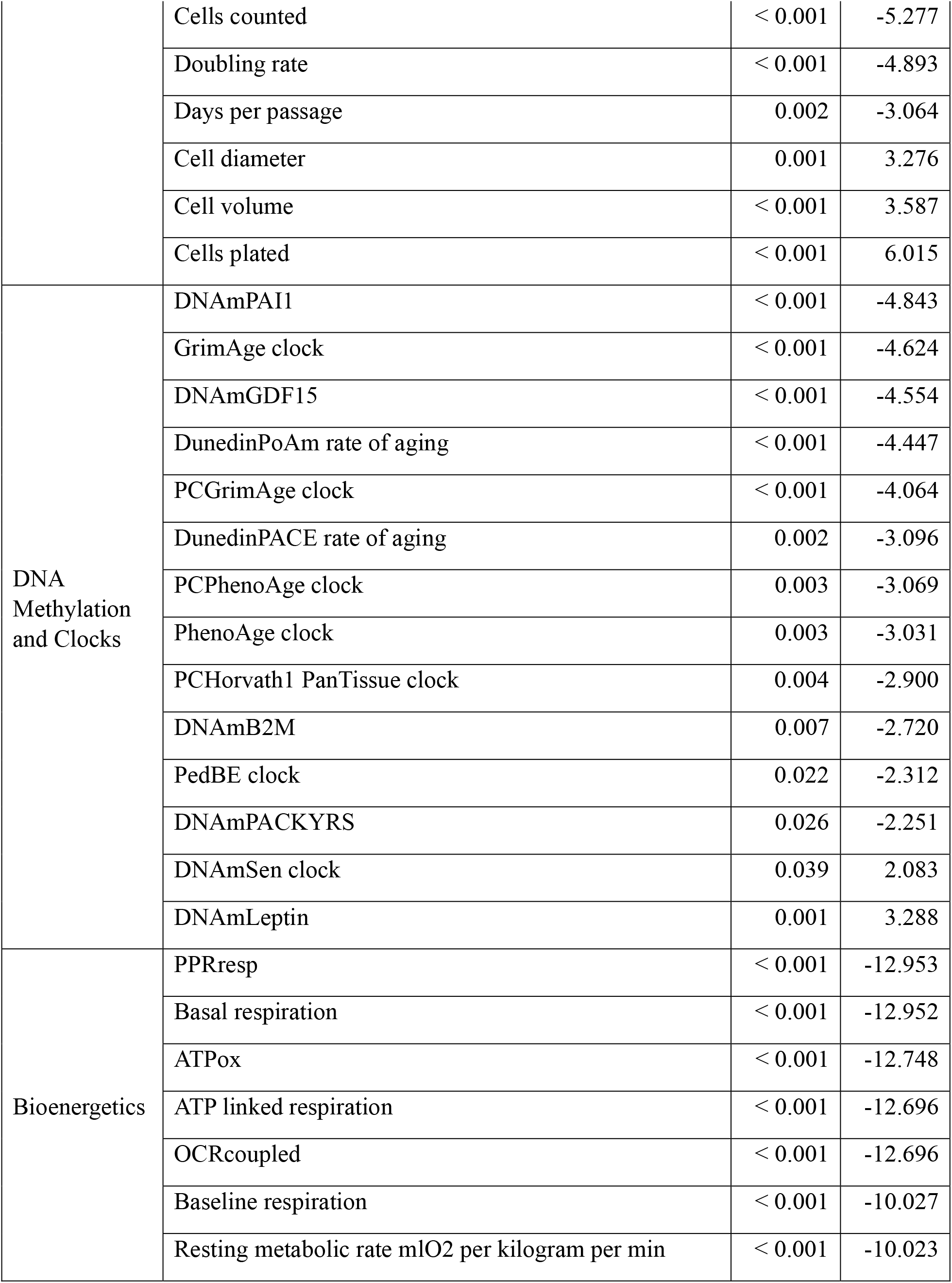
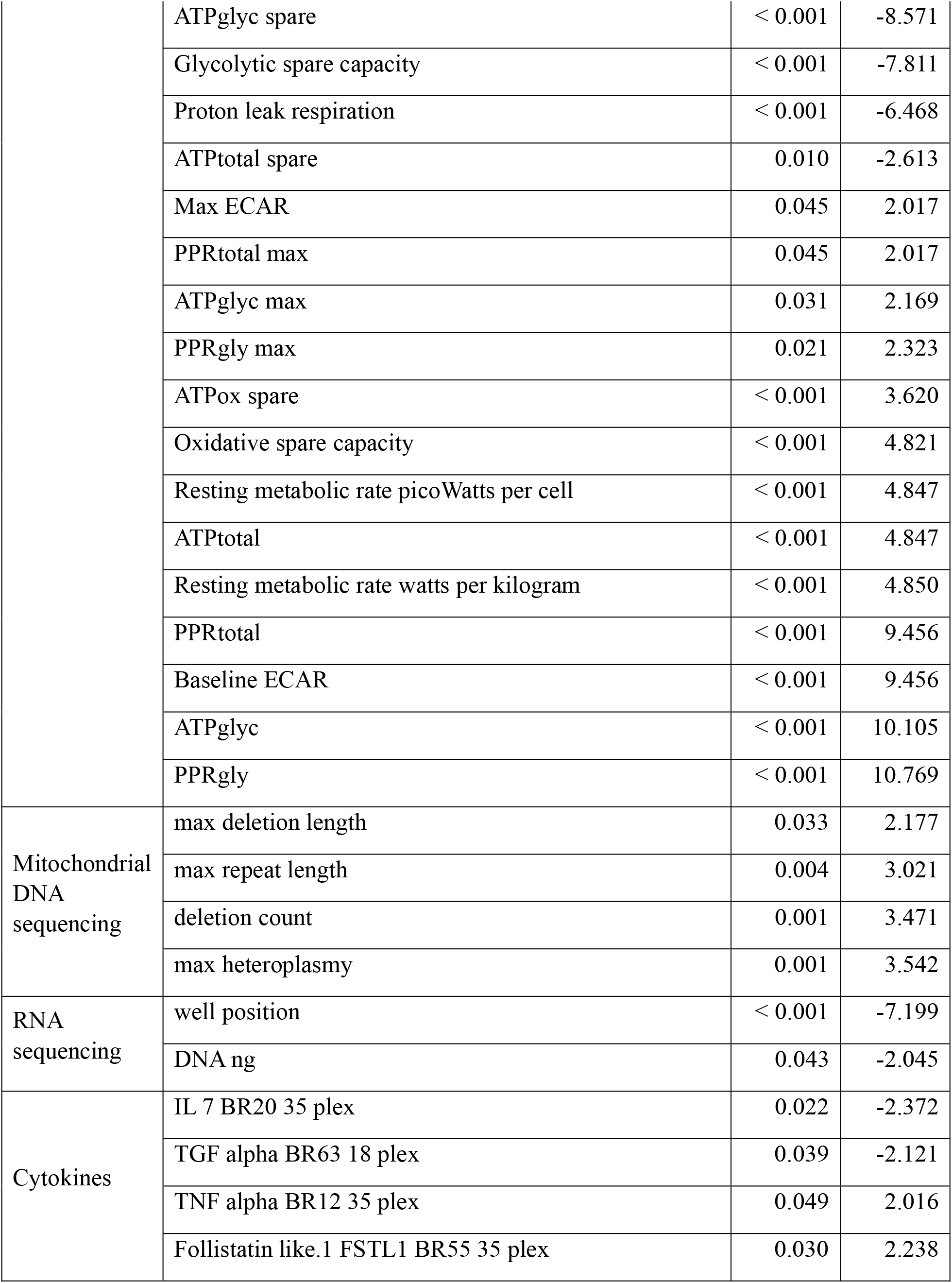
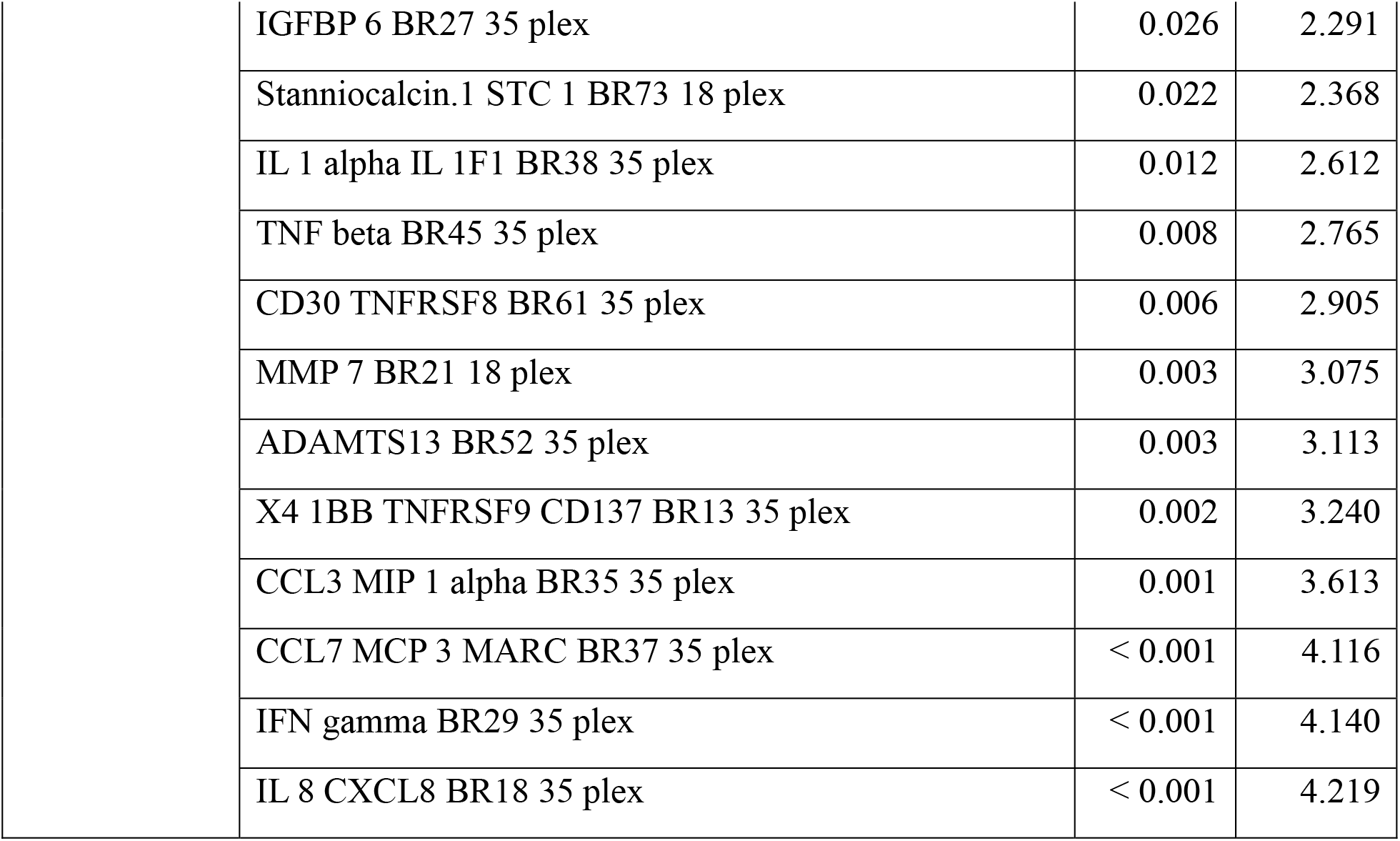
Quantifying impact of SURF1 mutation on biomarkers.

| Category | Biomarker | p-value | t-value |
| --- | --- | --- | --- |
| Cytology | Population doublings | < 0.001 | -9.659 |
|  | Divisions per passage | < 0.001 | -6.754 |
|  | Live cells counted | < 0.001 | -5.291 |
|  | Cells counted | < 0.001 | -5.277 |
|  | Doubling rate | < 0.001 | -4.893 |
|  | Days per passage | 0.002 | -3.064 |
|  | Cell diameter | 0.001 | 3.276 |
|  | Cell volume | < 0.001 | 3.587 |
|  | Cells plated | < 0.001 | 6.015 |
| DNA Methylation and Clocks | DNAmPAI1 | < 0.001 | -4.843 |
|  | GrimAge clock | < 0.001 | -4.624 |
|  | DNAmGDF15 | < 0.001 | -4.554 |
|  | DunedinPoAm rate of aging | < 0.001 | -4.447 |
|  | PCGrimAge clock | < 0.001 | -4.064 |
|  | DunedinPACE rate of aging | 0.002 | -3.096 |
|  | PCPhenoAge clock | 0.003 | -3.069 |
|  | PhenoAge clock | 0.003 | -3.031 |
|  | PCHorvath1 PanTissue clock | 0.004 | -2.900 |
|  | DNAmB2M | 0.007 | -2.720 |
|  | PedBE clock | 0.022 | -2.312 |
|  | DNAmPACKYRS | 0.026 | -2.251 |
|  | DNAmSen clock | 0.039 | 2.083 |
|  | DNAmLeptin | 0.001 | 3.288 |
| Bioenergetics | PPRresp | < 0.001 | -12.953 |
|  | Basal respiration | < 0.001 | -12.952 |
|  | ATPox | < 0.001 | -12.748 |
|  | ATP linked respiration | < 0.001 | -12.696 |
|  | OCRCoupled | < 0.001 | -12.696 |
|  | Baseline respiration | < 0.001 | -10.027 |
|  | Resting metabolic rate mlO2 per kilogram per min | < 0.001 | -10.023 |
|  | ATPglyc spare | < 0.001 | -8.571 |
|  | Glycolytic spare capacity | < 0.001 | -7.811 |
|  | Proton leak respiration | < 0.001 | -6.468 |
|  | ATPtotal spare | 0.010 | -2.613 |
|  | Max ECAR | 0.045 | 2.017 |
|  | PPRtotal max | 0.045 | 2.017 |
|  | ATPglyc max | 0.031 | 2.169 |
|  | PPRgly max | 0.021 | 2.323 |
|  | ATPox spare | < 0.001 | 3.620 |
|  | Oxidative spare capacity | < 0.001 | 4.821 |
|  | Resting metabolic rate picoWatts per cell | < 0.001 | 4.847 |
|  | ATPtotal | < 0.001 | 4.847 |
|  | Resting metabolic rate watts per kilogram | < 0.001 | 4.850 |
|  | PPRtotal | < 0.001 | 9.456 |
|  | Baseline ECAR | < 0.001 | 9.456 |
|  | ATPglyc | < 0.001 | 10.105 |
|  | PPRgly | < 0.001 | 10.769 |
| Mitochondrial DNA sequencing | max deletion length | 0.033 | 2.177 |
|  | max repeat length | 0.004 | 3.021 |
|  | deletion count | 0.001 | 3.471 |
|  | max heteroplasmy | 0.001 | 3.542 |
| RNA sequencing | well position | < 0.001 | -7.199 |
|  | DNA ng | 0.043 | -2.045 |
| Cytokines | IL 7 BR20 35 plex | 0.022 | -2.372 |
|  | TGF alpha BR63 18 plex | 0.039 | -2.121 |
|  | TNF alpha BR12 35 plex | 0.049 | 2.016 |
|  | Follistatin like.1 FSTL1 BR55 35 plex | 0.030 | 2.238 |

|  |  |  |
| --- | --- | --- |
| IGFBP 6 BR27 35 plex | 0.026 | 2.291 |
| Stanniocalcin.1 STC 1 BR73 18 plex | 0.022 | 2.368 |
| IL 1 alpha IL 1F1 BR38 35 plex | 0.012 | 2.612 |
| TNF beta BR45 35 plex | 0.008 | 2.765 |
| CD30 TNFRSF8 BR61 35 plex | 0.006 | 2.905 |
| MMP 7 BR21 18 plex | 0.003 | 3.075 |
| ADAMTS13 BR52 35 plex | 0.003 | 3.113 |
| X4 1BB TNFRSF9 CD137 BR13 35 plex | 0.002 | 3.240 |
| CCL3 MIP 1 alpha BR35 35 plex | 0.001 | 3.613 |
| CCL7 MCP 3 MARC BR37 35 plex | < 0.001 | 4.116 |
| IFN gamma BR29 35 plex | < 0.001 | 4.140 |
| IL 8 CXCL8 BR18 35 plex | < 0.001 | 4.219 |

Among all of the dependent variables (biomarkers), PPRgly, ATPglyc, Baseline ECAR, PPRtotal, and cells plated were the most positively impacted by the SURF1 mutation. PPRresp, basal respiration, ATPox, ATP linked respiration, OCRcoupled, baseline respiration, resting metabolic rate mlO2 per kilogram per min, and population doublings were the most negatively impacted by the SURF1 mutation. In addition, all of these dependent variables (biomarkers) listed above had p-values of less than 1e-6. The full results are thoroughly analyzed in the discussion section.

The final data table contains the result of student’s t-test for paired samples between the dependent variables (biomarkers) and experimental treatment (control, Dexamethasone, Oligomycin).

**Table 3:**
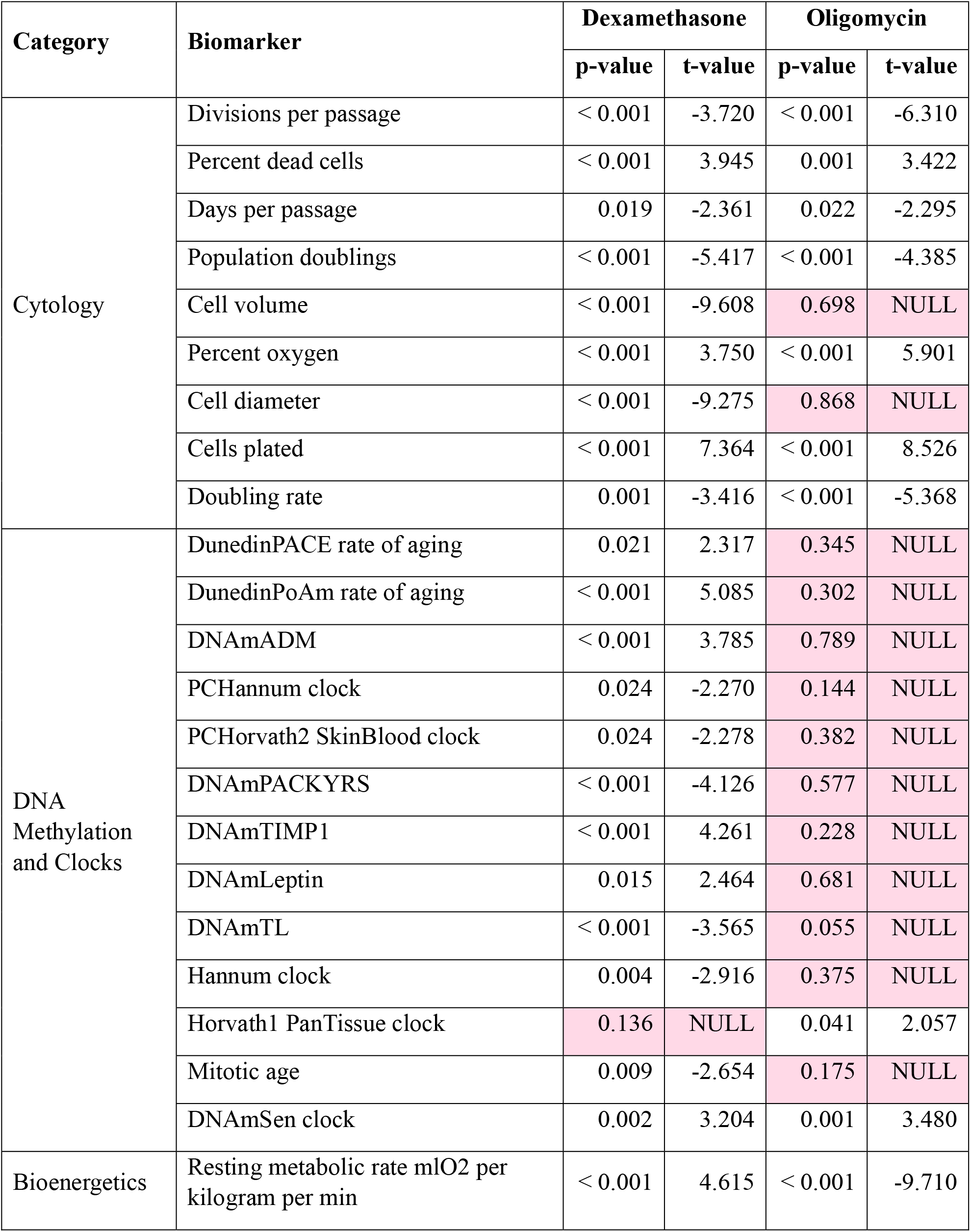
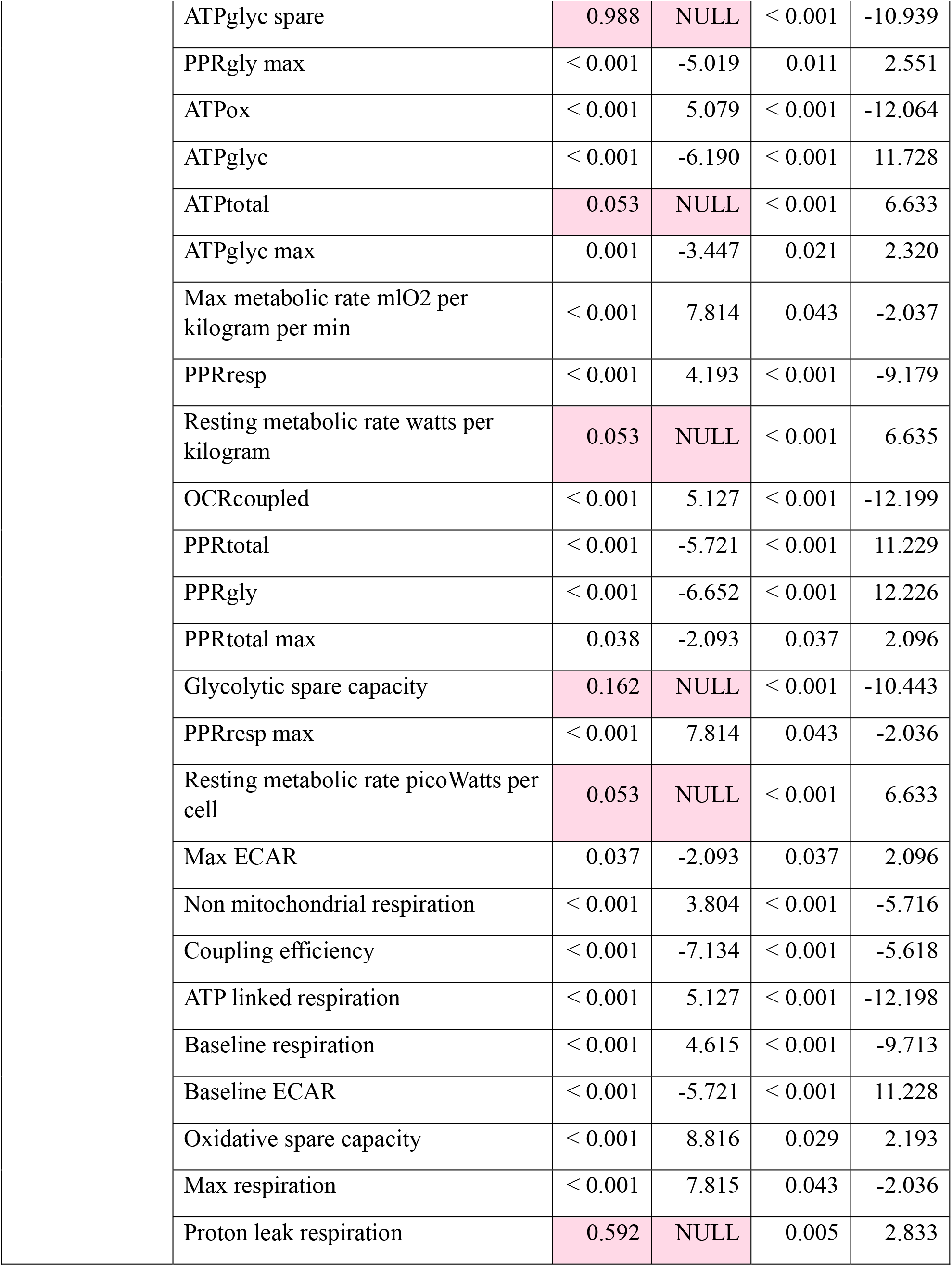
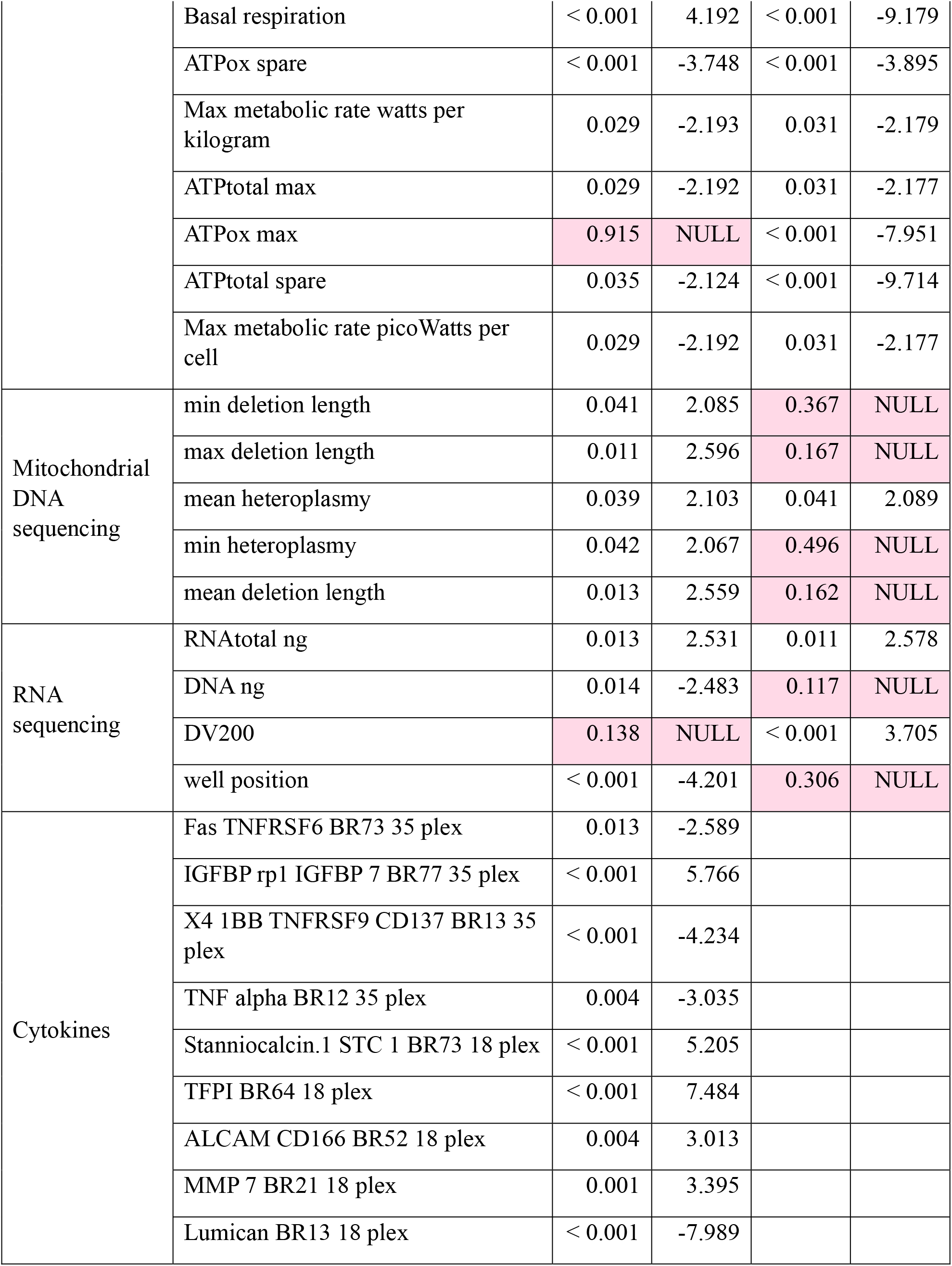
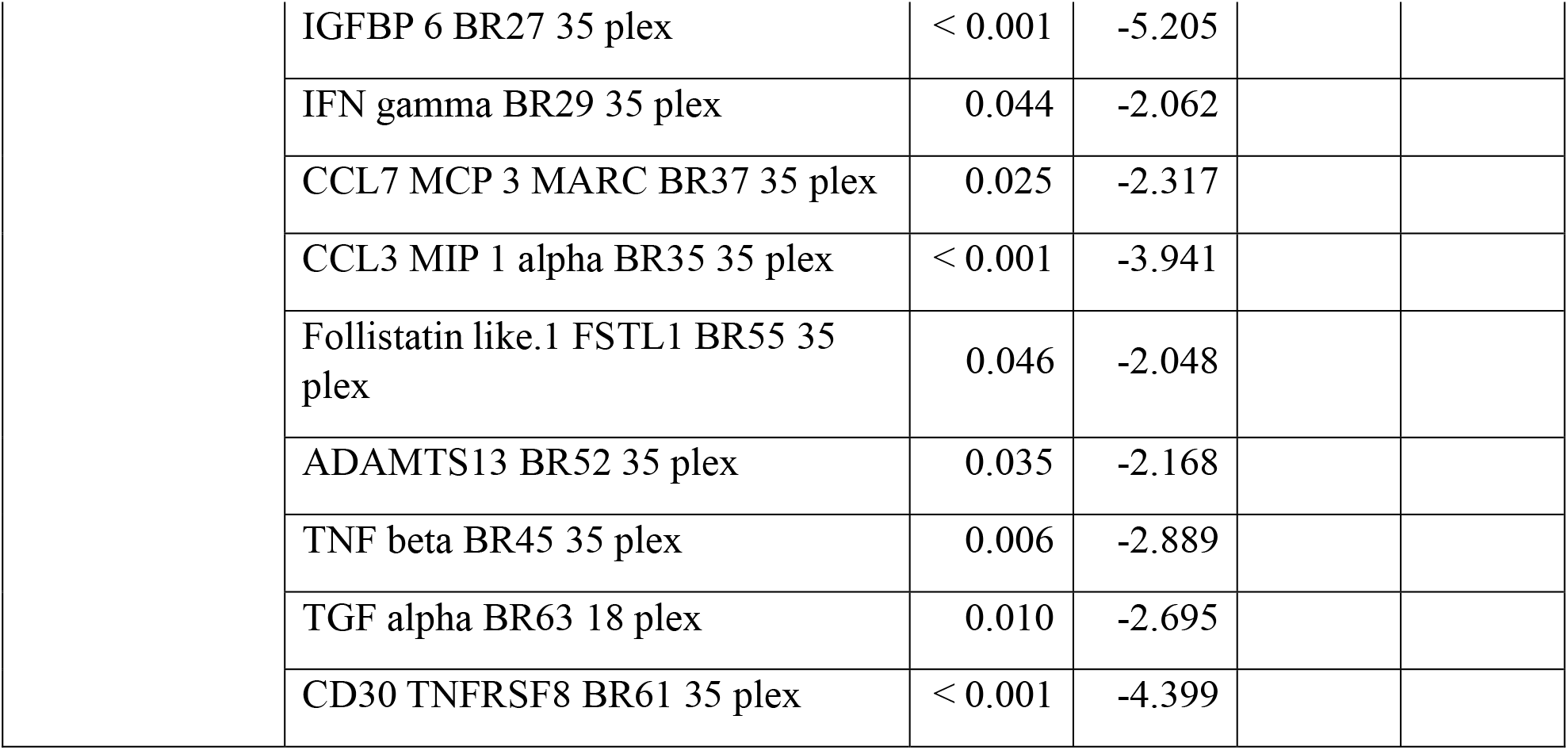
Quantifying impact of Dexamethasone and Oligomycin on biomarkers.

| Category | Biomarker | Dexamethasone |  | Oligomycin |  |
| --- | --- | --- | --- | --- | --- |
|  |  | p-value | t-value | p-value | t-value |
| Cytology | Divisions per passage | < 0.001 | -3.720 | < 0.001 | -6.310 |
|  | Percent dead cells | < 0.001 | 3.945 | 0.001 | 3.422 |
|  | Days per passage | 0.019 | -2.361 | 0.022 | -2.295 |
|  | Population doublings | < 0.001 | -5.417 | < 0.001 | -4.385 |
|  | Cell volume | < 0.001 | -9.608 | 0.698 | NULL |
|  | Percent oxygen | < 0.001 | 3.750 | < 0.001 | 5.901 |
|  | Cell diameter | < 0.001 | -9.275 | 0.868 | NULL |
|  | Cells plated | < 0.001 | 7.364 | < 0.001 | 8.526 |
|  | Doubling rate | 0.001 | -3.416 | < 0.001 | -5.368 |
| DNA Methylation and Clocks | DunedinPACE rate of aging | 0.021 | 2.317 | 0.345 | NULL |
|  | DunedinPoAm rate of aging | < 0.001 | 5.085 | 0.302 | NULL |
|  | DNAmADM | < 0.001 | 3.785 | 0.789 | NULL |
|  | PCHannum clock | 0.024 | -2.270 | 0.144 | NULL |
|  | PCHorvath2 SkinBlood clock | 0.024 | -2.278 | 0.382 | NULL |
|  | DNAmPACKYRS | < 0.001 | -4.126 | 0.577 | NULL |
|  | DNAmTIMP1 | < 0.001 | 4.261 | 0.228 | NULL |
|  | DNAmLeptin | 0.015 | 2.464 | 0.681 | NULL |
|  | DNAmTL | < 0.001 | -3.565 | 0.055 | NULL |
|  | Hannum clock | 0.004 | -2.916 | 0.375 | NULL |
|  | Horvath1 PanTissue clock | 0.136 | NULL | 0.041 | 2.057 |
|  | Mitotic age | 0.009 | -2.654 | 0.175 | NULL |
|  | DNAmSen clock | 0.002 | 3.204 | 0.001 | 3.480 |
| Bioenergetics | Resting metabolic rate mlO2 per kilogram per min | < 0.001 | 4.615 | < 0.001 | -9.710 |

|  |  |  |  |  |
| --- | --- | --- | --- | --- |
| ATPglyc spare | 0.988 | NULL | < 0.001 | -10.939 |
| PPRgly max | < 0.001 | -5.019 | 0.011 | 2.551 |
| ATPox | < 0.001 | 5.079 | < 0.001 | -12.064 |
| ATPglyc | < 0.001 | -6.190 | < 0.001 | 11.728 |
| ATPtotal | 0.053 | NULL | < 0.001 | 6.633 |
| ATPglyc max | 0.001 | -3.447 | 0.021 | 2.320 |
| Max metabolic rate mlO2 per kilogram per min | < 0.001 | 7.814 | 0.043 | -2.037 |
| PPRresp | < 0.001 | 4.193 | < 0.001 | -9.179 |
| Resting metabolic rate watts per kilogram | 0.053 | NULL | < 0.001 | 6.635 |
| OCRcoupled | < 0.001 | 5.127 | < 0.001 | -12.199 |
| PPRtotal | < 0.001 | -5.721 | < 0.001 | 11.229 |
| PPRgly | < 0.001 | -6.652 | < 0.001 | 12.226 |
| PPRtotal max | 0.038 | -2.093 | 0.037 | 2.096 |
| Glycolytic spare capacity | 0.162 | NULL | < 0.001 | -10.443 |
| PPRresp max | < 0.001 | 7.814 | 0.043 | -2.036 |
| Resting metabolic rate picoWatts per cell | 0.053 | NULL | < 0.001 | 6.633 |
| Max ECAR | 0.037 | -2.093 | 0.037 | 2.096 |
| Non mitochondrial respiration | < 0.001 | 3.804 | < 0.001 | -5.716 |
| Coupling efficiency | < 0.001 | -7.134 | < 0.001 | -5.618 |
| ATP linked respiration | < 0.001 | 5.127 | < 0.001 | -12.198 |
| Baseline respiration | < 0.001 | 4.615 | < 0.001 | -9.713 |
| Baseline ECAR | < 0.001 | -5.721 | < 0.001 | 11.228 |
| Oxidative spare capacity | < 0.001 | 8.816 | 0.029 | 2.193 |
| Max respiration | < 0.001 | 7.815 | 0.043 | -2.036 |
| Proton leak respiration | 0.592 | NULL | 0.005 | 2.833 |
|  | Basal respiration | < 0.001 | 4.192 | < 0.001 | -9.179 |
|  | ATPox spare | < 0.001 | -3.748 | < 0.001 | -3.895 |
|  | Max metabolic rate watts per kilogram | 0.029 | -2.193 | 0.031 | -2.179 |
|  | ATPtotal max | 0.029 | -2.192 | 0.031 | -2.177 |
|  | ATPox max | 0.915 | NULL | < 0.001 | -7.951 |
|  | ATPtotal spare | 0.035 | -2.124 | < 0.001 | -9.714 |
|  | Max metabolic rate picoWatts per cell | 0.029 | -2.192 | 0.031 | -2.177 |
| Mitochondrial DNA sequencing | min deletion length | 0.041 | 2.085 | 0.367 | NULL |
|  | max deletion length | 0.011 | 2.596 | 0.167 | NULL |
|  | mean heteroplasmy | 0.039 | 2.103 | 0.041 | 2.089 |
|  | min heteroplasmy | 0.042 | 2.067 | 0.496 | NULL |
|  | mean deletion length | 0.013 | 2.559 | 0.162 | NULL |
| RNA sequencing | RNAtotal ng | 0.013 | 2.531 | 0.011 | 2.578 |
|  | DNA ng | 0.014 | -2.483 | 0.117 | NULL |
|  | DV200 | 0.138 | NULL | < 0.001 | 3.705 |
|  | well position | < 0.001 | -4.201 | 0.306 | NULL |
| Cytokines | Fas TNFRSF6 BR73 35 plex | 0.013 | -2.589 |  |  |
|  | IGFBP rp1 IGFBP 7 BR77 35 plex | < 0.001 | 5.766 |  |  |
|  | X4 1BB TNFRSF9 CD137 BR13 35 plex | < 0.001 | -4.234 |  |  |
|  | TNF alpha BR12 35 plex | 0.004 | -3.035 |  |  |
|  | Stanniocalcin.1 STC 1 BR73 18 plex | < 0.001 | 5.205 |  |  |
|  | TFPI BR64 18 plex | < 0.001 | 7.484 |  |  |
|  | ALCAM CD166 BR52 18 plex | 0.004 | 3.013 |  |  |
|  | MMP 7 BR21 18 plex | 0.001 | 3.395 |  |  |
|  | Lumican BR13 18 plex | < 0.001 | -7.989 |  |  |

|  |  |  |
| --- | --- | --- |
| IGFBP 6 BR27 35 plex | < 0.001 | -5.205 |
| IFN gamma BR29 35 plex | 0.044 | -2.062 |
| CCL7 MCP 3 MARC BR37 35 plex | 0.025 | -2.317 |
| CCL3 MIP 1 alpha BR35 35 plex | < 0.001 | -3.941 |
| Follistatin like.1 FSTL1 BR55 35 plex | 0.046 | -2.048 |
| ADAMTS13 BR52 35 plex | 0.035 | -2.168 |
| TNF beta BR45 35 plex | 0.006 | -2.889 |
| TGF alpha BR63 18 plex | 0.010 | -2.695 |
| CD30 TNFRSF8 BR61 35 plex | < 0.001 | -4.399 |

Among all of the dependent variables (biomarkers), Oxidative spare capacity, max respiration, max metabolic rate mlO2 per kilogram per min, PPRresp max, TFPI BR64 18 plex, cells plated, and IGFBP rp1 IGFBP 7 BR77 35 plex were the most positively impacted by Dexamethasone. Cell volume, cell diameter, Lumican BR13 18 plex, coupling efficiency, PPRgly, ATPglyc, PPRtotal, and baseline ECAR were the most negatively impacted by Dexamethasone. In addition, all of these dependent variables (biomarkers) listed above had p-values of less than 1e-6. The full results are thoroughly analyzed in the discussion section.

The Columbia University researchers did not measure cytokines in normal fibroblasts with Oligomycin as the experimental treatment, which is why there is a blank section in the data table above. Among all of the dependent variables (biomarkers), PPRgly, ATPglyc, PPRtotal, baseline ECAR, cells plated, resting metabolic rate watts per kilogram, ATPtotal, and resting metabolic rate picoWatts per cell were the most positively impacted by Oligomycin.

OCRcoupled, ATP linked respiration, ATPox, ATPglyc spare, glycolytic spare capacity, ATPtotal spare, baseline respiration, and resting metabolic rate mlO2 per kilogram per min were the mos negatively impacted by Oligomycin. In addition, all of these dependent variables (biomarkers) listed above had p-values of less than 1e-6. The full results are thoroughly analyzed in the discussion section.

## Discussion

In this section, we analyzed our findings across all six molecular and cellular data types we studied. For each category, we discussed correlations with biological age, the effects of the SURF1 mutation, and the influence of treatments like Dexamethasone and Oligomycin.

### Cytology

The doubling rate, population doubling time, population doublings, divisions per passage, and days per passage all measure a specific aspect of cell growth and proliferation. A doubling is when the population of cells has doubled, and a passage refers to the process of moving the cells to a new area containing fresh growth medium. Population doublings clearly had a positive correlation with age as it only ever increased over time. Days per passage was held constant, yet divisions per passage had a negative correlation with age. This provides evidence for the Hayflick limit, which states that cells undergo mitosis less frequently as they age, eventually stopping proliferation completely [17]. This is also supported by the fact that doubling rate also had a negative correlation with age. Population doublings, divisions per passage, and doubling rate all decreased in fibroblasts with the SURF1 mutation. It can be inferred that the energy deficiency slowed down the cell cycle. This explains why individuals with Leigh syndrome experience weight faltering [18]. As expected, both Dexamethasone and Oligomycin negatively impacted these measurements as mitochondrial dysfunction and oxidative stress do not help with cell proliferation.

Live cells counted and percent dead cells refer to the amount of viable and nonviable cells in the sample. Live cells counted had a negative correlation with age while percent dead cells had a positive correlation with age. This is simply due to the Hayflick limit. The SURF1 mutation decreased live cells counted while Dexamethasone and Oligomycin increased the percent dead cells. This shows the direct impact of mitochondrial dysfunction and oxidative stress on the cell cycle [19]. Finally, cell diameter and volume had a negative correlation with age. This could have been caused by most cells being in the S, G2, or M phase at the start of the experiment while spending more time in G0 or G1 towards the end of the experiment. Finally, cell diameter and volume were increased by the SURF1 mutation. This was most likely caused by impaired mitochondria being unable to maintain ion homeostasis with the environment, causing osmotic swelling [20].

### DNA Methylation and Clocks

All of the dependent variables ending with the word “clock” are epigenetic clocks that measure the biological age of the fibroblasts. When there is a “PC” before the name of the epigenetic clock, it refers to the principal component of the clock. The Hannum and PhenoAge clocks along with their principal components had a strong positive correlation with total days grown. This validates the experimental setup as the fibroblasts were indeed aging properly in the cell culture. GrimAge and PhenoAge along with their principal components were lower in cells with the SURF1 mutation. This is easily explained by the fact that fibroblasts with this mutation died after less than 150 days while some healthy ones survived over 500 days. Dexamethasone negatively impacted Hannum clock and its principal component. This was probably caused by a suppressed immune system and modified glucose metabolism, causing more DNA methylation [21]. The fact that Horvath1 PanTissue clock and PCHorvath2 SkinBlood clock also were negatively impacted by Dexamethasone supports the emerging idea that NDMI could be used as a biomarker for evaluation glucocorticoid treatments [22]. DunedinPACE and DunedinPoAm both had a negative correlation with biological age and Dexamethasone. This supports the Reliability Theory of Aging and Longevity which states that as cells age, the rate of aging slows [23].

DNAmLeptin quantifies the amount of methylation of the *LEP* gene promoter. It had a positive correlation with age, SURF1 mutation, and Dexamethasone. It follows that as humans age, leptin levels are reduced, impacting energy expenditure and appetite regulation [24]. Mitotic age estimates the total number of cell divisions that occurred. This trivially explains its strong positive correlation with age. DNAmPAI1 quantifies the amount of methylation on the *SERPINE1* gene that encodes plasminogen activator inhibitor-1 (PAI1). Its negative correlation with age provides a biological explanation of why cancer and blot clots are more prevalent in older populations. This is because PAI1 has been shown to increase thrombosis and found in extreme amounts in biopsies of tumors [25]. SURF1 negatively impacting DNAmPAI1 also hints at a correlation between mitochondrial dysfunction and risk of cancer. DNAmB2M refers to amount of methylation on the *B2M* gene, which encodes β_2_ microglobulin. It also had a negative correlation with age. Elevated amounts of this MHC class I molecule have been linked to kidney problems; this provides an explanation for why kidney diseases are more prevalent in the elderly [26]. β_2_ microglobulin being negatively impacted by SURF1 mutation is due to its correlation with oxidative stress [27]. DNAmADM quantifies the amount of methylation on the *ADM* gene, which codes for Adrenomedullin, which is a vasodilator peptide hormone. It had a negative correlation with age, meaning that more Adrenomedullin is produced in elderly, which is a risk factor for Alzheimer’s disease [28].

### Bioenergetics

First, we will investigate variables related to glycolysis and oxidative phosphorylation.

Most of these such as PPRgly, ATPglyc, Baseline ECAR, PPRresp max, etc. had a positive correlation with age. This indicates that as fibroblasts age, they require more energy to maintain homeostasis through stress or damage repair. Plus, a strong correlation with the glycolysis variables indicates a heightened reliance on glycolysis as cells age. This is due to an increased demand for rapid ATP production. Oxidative phosphorylation was also increased in older cells. This suggests an alternative theory that there is metabolic inefficiency as cells age or mitochondrial dysregulation [29]. Coupling efficiency had a negative correlation with aging, which implies that mitochondria in the elderly have higher proton leak or ROS, which supports the free radical theory of aging. The SURF1 mutation led to higher values for glycolysis statistics and lower values for oxidative phosphorylation. This is a trivial result of COX deficiency due to a mutation in the *SURF1* gene. Interestingly, resting metabolic rate was decreased due to Oligomycin. This is most likely due to a reduction in respiration rate to maintain a proton gradient and a shift towards glycolysis [30].

### Mitochondrial DNA Sequencing

All of the deletion columns refer to deletions in the mtDNA sequence, where one or more base pairs are absent when compared to the reference genome. None of them were correlated with biological age or impacted by Oligomycin, but some deletion columns were impacted by SURF1 mutation and Dexamethasone. Deletion count is the total number of deletions and was positively impacted by the SURF1 mutation. A high number of deletions causes genomic instability, leading to mitochondrial dysfunction such as Kearns-Sayre syndrome [31]. Mean deletion length refers to the average size of all the deletions. The mean deletion length increased due to Dexamethasone, which confirms that this corticosteroid can disrupt mtDNA’s function in patients [32]. Max deletion length had a positive correlation with SURF1 while min deletion length had no correlation. This also supports the theory that the SURF1 mutation indirectly leads to genomic instability. The fact that both max and min deletion length increased due to Dexamethasone points to the fact that this glucocorticoid agonist disrupts mtDNA repair through stress-mediated pathways.

All of the repeat columns refer to repeat sequences in the mtDNA which are caused by a variety of biological processes such as polymerase slippage [33]. Max repeat length had a negative correlation with age. This is probably because long repeat sequences lead to secondary structures such as hairpins, which are more susceptible to deletions [34]. These short repeat sequences could lead to mitochondrial dysfunction; this provides a biological explanation of why diseases such as Alzheimer’s, Parkinson’s, and Amyotrophic Lateral Sclerosis are correlated with age [35]. The positive impact of the SURF1 mutation on max repeat length is because of impaired COX, leading to lower ROS. This slows the rate of oxidative damage to mtDNA, leading to a higher max repeat length [36]. This supports the free radical theory of aging. Finally, no repeat columns were impacted by Dexamethasone or Oligomycin.

All of the heteroplasmy columns refer to the proportion of mtDNA molecules that have a mutation. In humans, each mutation found in about 1 to 2% of all mtDNA is referred to as microheteroplasmy. Max heteroplasmy was higher in fibroblasts with the SURF1 mutation. This suggests that mtDNA with mutations may persist longer in mitochondria undergoing oxidative stress. Both mean and min heteroplasmy were increased in fibroblasts cultured with Dexamethasone. This implies that Dexamethasone may reduce selection in mutated mtDNA through increased ROS and fewer fusion/fission events, leading to impaired mitophagy causing aerobic glycolysis to be the primary ATP source [37]. Mitochondrial dysfunction due to aging could create a positive feedback loop of mtDNA mutations accumulating, eventually causing cancer due to the Warburg effect [38]. Oligomycin positively impacting mean heteroplasmy further supports this argument and the free radical theory of aging.

### RNA Sequencing

DV200 is the percent of RNA fragments that were greater than 200 nucleotides long. Although its primary purpose is to quantify RNA fragmentation, it also provides some insight into the original RNA condition in cells. DV200’s negative correlation with age implies that oxidative stress along with RNA turnover due to changes in ribonuclease pathways are hallmarks of aging [39, 40]. DV200 being higher in fibroblasts treated with Oligomycin further supports the theory that oxidative stress contributes to RNA degradation in fibroblasts [41]. RNA total ng and DNA ng refer to the amount of RNA and DNA contamination in the extracted sample in nanograms. DNA ng was negatively impacted by the SURF1 mutation because the stress inhibited proliferation and disrupts the cell cycle [41]. Dexamethasone upregulates some genes for anti-inflammatory responses, which explains why RNA total ng increased in fibroblasts treated with it [42]. In contrast, DNA ng decreased because Dexamethasone inhibited the cell cycle, disrupting the synthesis of new DNA strands. Similarly, Oligomycin increased RNA total ng as the cells upregulated RNA transcription as a response to its metabolic stress. Well position refers to the specific index on a well plate and its negative impact from SURF1 and Dexamethasone are merely due to the researchers’ well location choice. RIN refers to the RNA integrity number that quantifies the quality of RNA through electrophoresis. RIN had a negative correlation with age, and this was probably the result of RNA degradation over time [43].

Finally, the negative correlation Base 30 and quality had with age was due to a few outliers.

### Cytokines

First, we will start with the interleukins and other miscellaneous cytokines. IL18, also known as IL1F4 or interferon-gamma inducing factor, is a proinflammatory protein that in excessive amounts has been linked to Alzheimer’s and heart failure [44, 45]. Therefore, its positive correlation with age hints that a weaker blood-brain barrier coupled with Amyloid-β accumulation could cause Alzheimer’s. Heart failure is most likely caused by increased production of interferon-γ by T cells, leading to plaque formation in blood vessels. IL1 alpha, also known as hematopoietin 1, is activated by the immune system in humans to induce fever and inflammatory responses. PCSK9, also known as proprotein convertase subtilisin/kexin type 9, is an enzyme highly expressed in the skin for cholesterol and free fatty acid synthesis. It had a negative correlation with age, indicating that older people are at higher risk of coronary artery disease due to LDL receptor degradation [46]. TFPI, also known as tissue factor pathway inhibitor, is a single-chain protein that prevents blood coagulation and regulates clotting. Its negative correlation with age explains why older people are more susceptible to ischemic stroke; a lack of TFPI enhances coagulation pathways which directly causes vascular complications [47].

CXCL8, also known as IL8 or C-X-C motif chemokine ligand 8, is both a chemokine and interleukin produced by epithelial cells to attract neutrophils to infection sites. Its negative correlation with age means that older people would have reduced inflammation at the cost of less effective immune responses. This provides a biological explanation for why the elderly have a harder time fighting pathogens [48]. CCL7, also known as monocyte-chemotactic protein 3, is a chemoattractant for leukocytes in the immune system. It has a positive correlation with aging, increasing risk of multiple sclerosis due to leukocyte migration across the blood-brain barrier [49]. The amount of CCL7 was also increased by the SURF1 mutation, hinting at link between mitochondrial dysfunction and inflammatory immune response most likely caused by a high amount of ROS.

TGF alpha is a ligand that can start the EGFR pathway and is involved in tissue repair. Its positive correlation with age is most likely due to cellular senescence. GDF15 stands for growth/differentiation factor 15 and its positive correlation with age helps explain that skin thins in elderly people as mitochondrial dysfunction upregulates this protein [50]. Insulin-like growth factor-binding protein 6 (IGFBP6) is a transport protein for IGF-1, which activates the Akt signaling pathway. There was a negative correlation between IGFBP6 and age. This explains the mechanism behind why cells undergo mitosis less often as they age, which was analyzed in the cytology section.

Tumor necrosis factor receptor superfamily (TNFRSF) refers to cytokine receptors that bind to TNFs and are usually found in the plasma membrane of cells. TNFRSF6 (Fas), TNFRSF8 (CD30), and TNFRSF9 (4-1BB) all had a strong negative correlation with age. Fas directly leads to programmed cell death, so lower levels of it could lead to cancer if there is not enough of it to kill cancerous cells. CD30 activates B and T cells and has been shown to protect the human body against autoimmunity. Therefore, it follows that autoimmunity is more common in older people, which precisely matches clinical studies [51]. Finally, 4-1BB stimulates T-cell proliferation so a decrease in this type 1 transmembrane protein impairs the ability of the elderly to have an effective adaptive immune response. TNF alpha and beta were negatively impacted by Dexamethasone due to its anti-inflammatory effects and the inhibition of NF-κB activation [52].

## Conclusion

Our research shed light on the underlying biochemical mechanisms of aging, shaped by many epigenetic and environmental factors. Analysis of the high temporal resolution multi-omics dataset revealed several aging biomarkers across various molecular and cellular data types. It also provided insights into how various biological pathways are affected. For example, increased population doublings and percent dead cells reflected cellular aging and degeneration in cytology. In mitochondrial DNA sequencing, positive aging effects included higher deletion counts and heteroplasmy, whereas negative aspects like shortened repeat lengths increased vulnerability to age-related disorders. In RNA sequencing, increased RNA total indicated stress adaptation. On the other hand, RNA degradation and lower RIN scores affected protein synthesis.

Our analysis demonstrated the importance of cytokines in aging-related diseases. Among the inflammatory cytokines, the increased levels of IL18, IL1 alpha, and CCL7 can lead to Alzheimer’s disease, chronic inflammatory disorders, multiple sclerosis, and chronic tissue damage. When it comes to tissue degeneration, increased levels of TGF alpha cause cellular proliferation and abnormal tissue repair that can lead to fibrosis and cancer; elevated levels of GDF15 correlate with reduced tissue integrity in elderly populations; whereas the reduced level of IGFBP6 impairs cell division, contributing to poor wound healing and tissue aging.

Additionally, as cells age, imbalances in certain cytokines affect the immune system. For example, reduced levels of TNFRSF6 increase cancer risks and failure to control autoimmune diseases; low levels of TNFRSF8 cause autoimmune disorders in the elderly; and aging-associated decreases in TNFRSF9 cause weakened adaptive immunity. Finally, coagulation and cardiovascular risks increase with decreased PCSK9 and TFPI activity in aging. These findings demonstrate the complex cytokine networks in aging and present potential targets for precision medicine in age-related health issues.

By comparing cells with the SURF1 mitochondrial mutation, associated with Leigh syndrome and Charcot–Marie–Tooth disease, to healthy ones, we had many insights into mitochondrial dysfunction and oxidative stress as accelerants of cellular senescence, supporting the free radical theory of aging while providing novel molecular analyses. The SURF1 mutation decreased doubling rates, population doublings, and divisions per passage. This resulted in impaired tissue regeneration and increased cellular senescence. SURF1 mutation decreased the live cell count highlighting accelerated cell death due to mitochondrial dysfunction. In cells with SURF1 mutation, an increase in cell diameter and volume due to osmotic swelling led to structural instability. Finally, the mutation lowered epigenetic clocks, GrimAge and PhenoAge, suggesting premature cellular aging.

We analyzed fibroblasts cultured in media containing Dexamethasone and Oligomycin. Dexamethasone reduced inflammation indicating its potential use in treating autoimmune diseases and allergies. However, it increased susceptibility to infections and slower tissue recovery. Additionally, it increased DNA methylation, reduced mitochondrial efficiency, lowered Hannum clock values, and reduced cell cycle activity that could lead to increased risk of chronic conditions, mitochondrial dysfunction, reduced cellular lifespan, and delayed healing across tissues. Oligomycin increased RNA response to stress enhancing transcription to rapidly address metabolic stress. This helped in short-term cellular survival. On the other hand, Oligomycin hampered mitochondrial ATP production causing energy deficits in high-demand tissues; increased glycolysis leading to cellular stress and metabolic imbalance; reduced basal metabolic respiration; and higher RNA degradation indicating oxidative stress-driven damage that contributed to reduced protein synthesis capabilities. Overall, Dexamethasone and Oligomycin treatments on fibroblasts showed negative impacts on their bioenergetics, mitochondrial function, and biological age in comparison with the control group without these treatments.

Our study provided insights into cellular aging that could be effective in prevention and treatment strategies for age-associated diseases. However, our study was limited to a specific cell type – fibroblasts. In the future, we plan to expand our research by including other cell types, a broader range of genetic mutations, and additional treatments. This approach will help develop a more comprehensive understanding of the aging process.

## Acknowledgements

Rajarshi Mandal thanks MIT PRIMES research program for giving him the opportunity to work with Professor Alterovitz on this project.

## References

[1] da Costa, J. P., Vitorino, R., Silva, G. M., Vogel, C., Duarte, A. C., & Rocha-Santos, T. (2016). A synopsis on aging—Theories, mechanisms and future prospects. Ageing Research Reviews, 29, 90–112. 10.1016/j.arr.2016.06.005

[2] Kirkwood, Thomas B. L., & Melov, S. (2011). On the Programmed/Non-Programmed Nature of Ageing within the Life History. Current Biology, 21(18), R701–R707. 10.1016/j.cub.2011.07.020

[3] Pomatto, L. C. D., & Davies, K. J. A. (2018). Adaptive homeostasis and the free radical theory of ageing. Free Radical Biology and Medicine, 124, 420–430. 10.1016/j.freeradbiomed.2018.06.016

[4] Lennicke, C., & Cochemé, H. M. (2020). Redox signalling and ageing: insights from Drosophila. Biochemical Society Transactions, 48(2), 367–377. 10.1042/bst20190052

[5] Ziada, A. S., Smith, M.-S. R., & Côté, H. C. F. (2020). Updating the Free Radical Theory of Aging. Frontiers in Cell and Developmental Biology, 8. 10.3389/fcell.2020.575645

[6] Dupont, C., Armant, D. R., & Brenner, C. (2009). Epigenetics: Definition, Mechanisms and Clinical Perspective. Seminars in Reproductive Medicine, 27(05), 351–357. 10.1055/s-0029-1237423

[7] Galupa, R., & Heard, E. (2018). X-Chromosome Inactivation: A Crossroads Between Chromosome Architecture and Gene Regulation. Annual Review of Genetics, 52(1), 535–566. 10.1146/annurev-genet-120116-024611

[8] Jansz, N. (2019). DNA methylation dynamics at transposable elements in mammals. Essays in Biochemistry. 10.1042/ebc20190039

[9] Duan, R., Fu, Q., Sun, Y., & Li, Q. (2022). Epigenetic clock: A promising biomarker and practical tool in aging. Ageing Research Reviews, 81, 101743. 10.1016/j.arr.2022.101743

[10] Sturm, G., Monzel, A. S., Karan, K. R., Michelson, J., Ware, S. A., Cardenas, A., Lin, J., Céline Bris, Santhanam, B., Murphy, M. P., Levine, M. E., Horvath, S., Belsky, D. W., Wang, S., Procaccio, V., Kaufman, B. A., Hirano, M., & Picard, M. (2022). A multi-omics longitudinal aging dataset in primary human fibroblasts with mitochondrial perturbations. Scientific Data, 9(1). 10.1038/s41597-022-01852-y

[11] Mick, David U., Dennerlein, S., Wiese, H., Reinhold, R., Pacheu-Grau, D., Lorenzi, I., Sasarman, F., Weraarpachai, W., Shoubridge, Eric A., Warscheid, B., & Rehling, P. (2012). MITRAC Links Mitochondrial Protein Translocation to Respiratory-Chain Assembly and Translational Regulation. Cell, 151(7), 1528–1541. 10.1016/j.cell.2012.11.053

[12] Echaniz-Laguna, A., Ghezzi, D., Chassagne, M., Mayencon, M., Padet, S., Melchionda, L., Rouvet, I., Lannes, B., Bozon, D., Latour, P., Zeviani, M., & Mousson de Camaret, B. (2013). SURF1 deficiency causes demyelinating Charcot-Marie-Tooth disease. Neurology, 81(17), 1523–1530. 10.1212/wnl.0b013e3182a4a518

[13] Huebner, K. D., Shrive, N. G., & Frank, C. B. (2013). Dexamethasone inhibits inflammation and cartilage damage in a new model of post-traumatic osteoarthritis. Journal of Orthopaedic Research, 32(4), 566–572. 10.1002/jor.22568

[14] Keeney, G. E., Gray, M. P., Morrison, A. K., Levas, M. N., Kessler, E. A., Hill, G. D., Gorelick, M. H., & Jackson, J. L. (2014). Dexamethasone for Acute Asthma Exacerbations in Children: A Meta-analysis. PEDIATRICS, 133(3), 493–499. 10.1542/peds.2013-2273

[15] Ahmed, M. H., & Hassan, A. (2020). Dexamethasone for the Treatment of Coronavirus Disease (COVID-19): a Review. SN Comprehensive Clinical Medicine, 2(12). 10.1007/s42399-020-00610-8

[16] Symersky, J., Osowski, D., Walters, D. E., & Mueller, D. M. (2012). Oligomycin frames a common drug-binding site in the ATP synthase. Proceedings of the National Academy of Sciences of the United States of America, 109(35), 13961–13965. 10.1073/pnas.1207912109

[17] Chan, M., Yuan, H., Soifer, I., Maile, T. M., Wang, R. Y., Ireland, A., O’Brien, J. J., Goudeau, J., Chan, L. J., Vijay, T., Freund, A., Kenyon, C., Bennett, B. D., McAllister, F. E., Kelley, D. R., Roy, M., Cohen, R. L., Levinson, A. D., Botstein, D., & Hendrickson, D. G. (2022). Novel insights from a multiomics dissection of the Hayflick limit. ELife, 11, e70283. 10.7554/eLife.70283

[18] Baskaran, D., & Hussain, N. (2020). Facial dysmorphism, hirsutism, and failure to thrive as manifestation of Leigh syndrome in a child with SURF1 mutation. Journal of Pediatric Neurosciences, 15(2), 108. 10.4103/jpn.jpn_137_18

[19] Byun, H.-O., Kim, H. Y., Lim, J. J., Seo, Y.-H., & Yoon, G. (2008). Mitochondrial dysfunction by complex II inhibition delays overall cell cycle progression via reactive oxygen species production. Journal of Cellular Biochemistry, 104(5), 1747–1759. 10.1002/jcb.21741

[20] Shimizu, T., Numata, T., & Okada, Y. (2004). A role of reactive oxygen species in apoptotic activation of volume-sensitive Cl-channel. Proceedings of the National Academy of Sciences of the United States of America, 101(17), 6770–6773. 10.1073/pnas.0401604101

[21] Morales-Nebreda, L., McLafferty, F. S., & Singer, B. D. (2019). DNA methylation as a transcriptional regulator of the immune system. Translational Research, 204, 1–18. 10.1016/j.trsl.2018.08.001

[22] Wiencke, J. K., Molinaro, A. M., Warrier, G., Rice, T., Clarke, J., Taylor, J., Wrensch, M., Hansen, H. M., McCoy, L., Tang, E., Tamaki, S. J., Tamaki, C. M., Nissen, E., Bracci, P., Salas, L. A., Koestler, D. C., Christensen, B. C., Zhang, Z., & Kelsey, K. T. (2022). DNA methylation as a pharmacodynamic marker of glucocorticoid response and glioma survival. 13(1). 10.1038/s41467-022-33215-x

[23] Gavrilov, L. A., & Gavrilova, N. S. (2005). Reliability Theory of Aging and Longevity. Elsevier EBooks, 3–42. 10.1016/b978-012088387-5/50004-2

[24] García-Cardona, M. C., Huang, F., García-Vivas, J. M., López-Camarillo, C., del Río Navarro, B. E., Navarro Olivos, E., Hong-Chong, E., Bolaños-Jiménez, F., & Marchat, L. A. (2014). DNA methylation of leptin and adiponectin promoters in children is reduced by the combined presence of obesity and insulin resistance. International Journal of Obesity, 38(11), 1457–1465. 10.1038/ijo.2014.30

[25] Li, S., Wei, X., He, J., Tian, X., Yuan, S., & Sun, L. (2018). Plasminogen activator inhibitor-1 in cancer research. Biomedicine & Pharmacotherapy, 105, 83–94. 10.1016/j.biopha.2018.05.119

[26] Argyropoulos, C. P., Chen, S. S., Ng, Y.-H., Roumelioti, M.-E., Shaffi, K., Singh, P. P., & Tzamaloukas, A. H. (2017). Rediscovering Beta-2 Microglobulin As a Biomarker across the Spectrum of Kidney Diseases. Frontiers in Medicine, 4. 10.3389/fmed.2017.00073

[27] Althubiti, M., Elzubier, M., Alotaibi, G. S., Althubaiti, M. A., Alsadi, H. H., Ziyad Abdulaziz Alhazmi, Alghamdi, F., Mahmoud Zaki El-Readi, Riyad Almaimani, & Abdullatif Babakr. (2021). Beta 2 microglobulin correlates with oxidative stress in elderly. Experimental Gerontology, 150, 111359–111359. 10.1016/j.exger.2021.111359

[28] Ferrero, H., Larrayoz, I. M., Martisova, E., Solas, M., Howlett, D. R., Francis, P. T., Gil-Bea, F. J., Martínez, A., & Ramírez, M. J. (2017). Increased Levels of Brain Adrenomedullin in the Neuropathology of Alzheimer’s Disease. Molecular Neurobiology, 55(6), 5177–5183. 10.1007/s12035-017-0700-6

[29] Baker, D. J., & Peleg, S. (2017). Biphasic Modeling of Mitochondrial Metabolism Dysregulation during Aging. Trends in Biochemical Sciences, 42(9), 702–711. 10.1016/j.tibs.2017.06.005

[30] Thoral, E., García-Díaz, C. C., Persson, E., Imen Chamkha, Eskil Elmér, Suvi Ruuskanen, & Nord, A. (2024). The relationship between mitochondrial respiration, resting metabolic rate and blood cell count in great tits. Biology Open, 13(3). 10.1242/bio.060302

[31] Comte, C., Tonin, Y., Heckel-Mager, A.-M., Boucheham, A., Smirnov, A., Auré, K., Lombès, A., Martin, R. P., Entelis, N., & Tarassov, I. (2013). Mitochondrial targeting of recombinant RNAs modulates the level of a heteroplasmic mutation in human mitochondrial DNA associated with Kearns Sayre Syndrome. Nucleic Acids Research, 41(1), 418–433. 10.1093/nar/gks965

[32] Luan, G., Li, G., Ma, X., Jin, Y., Hu, N., Li, J., Wang, Z., & Wang, H. (2019). Dexamethasone-Induced Mitochondrial Dysfunction and Insulin Resistance-Study in 3T3-L1 Adipocytes and Mitochondria Isolated from Mouse Liver. Molecules, 24(10), 1982. 10.3390/molecules24101982

[33] Zhang, H., Li, D., Zhao, X., Pan, S., Wu, X., Peng, S., Huang, H., Shi, R., & Tan, Z. (2020). Relatively semi-conservative replication and a folded slippage model for short tandem repeats. BMC Genomics, 21(1). 10.1186/s12864-020-06949-5

[34] Pan, F., Xu, P., Roland, C., Sagui, C., & Weninger, K. (2024). Structural and Dynamical Properties of Nucleic Acid Hairpins Implicated in Trinucleotide Repeat Expansion Diseases. Biomolecules, 14(10), 1278. 10.3390/biom14101278

[35] Somasundaram, I., Jain, S. M., Blot-Chabaud, M., Pathak, S., Banerjee, A., Rawat, S., Sharma, N. R., & Duttaroy, A. K. (2024). Mitochondrial dysfunction and its association with age-related disorders. Frontiers in Physiology, 15. 10.3389/fphys.2024.1384966

[36] Pharaoh, G., Pulliam, D., Hill, S., Sataranatarajan, K., & Van Remmen, H. (2016). Ablation of the mitochondrial complex IV assembly protein Surf1 leads to increased expression of the UPRMT and increased resistance to oxidative stress in primary cultures of fibroblasts. Redox Biology, 8, 430–438. 10.1016/j.redox.2016.05.001

[37] Wilasinee Suwanjang, Kay, Supaluk Prachayasittikul, Banthit Chetsawang, & Komgrid Charngkaew. (2019). Mitochondrial Dynamics Impairment in Dexamethasone-Treated Neuronal Cells. Neurochemical Research, 44(7), 1567–1581. 10.1007/s11064-019-02779-4

[38] Liberti, M. V., & Locasale, J. W. (2016). The Warburg Effect: How Does it Benefit Cancer Cells? Trends in Biochemical Sciences, 41(3), 211–218. 10.1016/j.tibs.2015.12.001

[39] Liu, M., Gong, X., Alluri, R. K., Wu, J., Sablo, T., & Li, Z. (2012). Characterization of RNA damage under oxidative stress in Escherichia coli. Bchm, 393(3), 123–132. 10.1515/hsz-2011-0247

[40] Tuck, A. C., & Tollervey, D. (2011). RNA in pieces. Trends in Genetics, 27(10), 422–432. 10.1016/j.tig.2011.06.001

[41] Dimozi, A., Mavrogonatou, E., Sklirou, A., & Kletsas, D. (2015). Oxidative stress inhibits the proliferation, induces premature senescence and promotes a catabolic phenotype in human nucleus pulposus intervertebral disc cells. European Cells and Materials, 30, 89–103. 10.22203/ecm.v030a07

[42] Giles, A. J., Hutchinson, M.-K. N. D., Sonnemann, H. M., Jung, J., Fecci, P. E., Ratnam, N. M., Zhang, W., Song, H., Bailey, R., Davis, D., Reid, C. M., Park, D. M., & Gilbert, M. R. (2018). Dexamethasone-induced immunosuppression: mechanisms and implications for immunotherapy. Journal for ImmunoTherapy of Cancer, 6(1). 10.1186/s40425-018-0371-5

[43] Wang, L., Nie, J., Sicotte, H., Li, Y., Eckel-Passow, J. E., Dasari, S., Vedell, P. T., Barman, P., Wang, L., Weinshiboum, R., Jen, J., Huang, H., Kohli, M., & Kocher, J.-P. A. (2016). Measure transcript integrity using RNA-seq data. BMC Bioinformatics, 17(1). 10.1186/s12859-016-0922-z

[44] Sutinen, E. M., Pirttilä, T., Anderson, G., Salminen, A., & Ojala, J. O. (2012). Pro-inflammatory interleukin-18 increases Alzheimer’s disease-associated amyloid-β production in human neuron-like cells. Journal of Neuroinflammation, 9(1). 10.1186/1742-2094-9-199

[45] O’Brien, L. C., Mezzaroma, E., Van Tassell, B. W., Marchetti, C., Carbone, S., Abbate, A., & Toldo, S. (2014). Interleukin-18 as a Therapeutic Target in Acute Myocardial Infarction and Heart Failure. Molecular Medicine, 20(1), 221–229. 10.2119/molmed.2014.00034

[46] Mega, J. L., Stitziel, N. O., Smith, J. G., Chasman, D. I., Caulfield, M. J., Devlin, J. J., Nordio, F., Hyde, C. L., Cannon, C. P., Sacks, F. M., Poulter, N. R., Sever, P. S., Ridker, P. M., Braunwald, E., Melander, O., Kathiresan, S., & Sabatine, M. S. (2015). Genetic risk, coronary heart disease events, and the clinical benefit of statin therapy: an analysis of primary and secondary prevention trials. The Lancet, 385(9984), 2264–2271. 10.1016/s0140-6736(14)61730-x

[47] Rossouw, J. E., Johnson, K. C., Pettinger, M., Cushman, M., Per Morten Sandset, Kuller, L., Frits Rosendaal, Rosing, J., Wasserthal-Smoller, S., Martin, L. W., Manson, J. E., Kamakshi Lakshminarayan, Merino, J. G., & Lynch, J. (2012). Tissue Factor Pathway Inhibitor, Activated Protein C Resistance, and Risk of Ischemic Stroke due to Postmenopausal Hormone Therapy. Stroke, 43(4), 952–957. 10.1161/strokeaha.111.643072

[48] Simon, A. K., Hollander, G. A., & McMichael, A. (2015). Evolution of the Immune System in Humans from Infancy to Old Age. Proceedings of the Royal Society B: Biological Sciences, 282(1821), 20143085. 10.1098/rspb.2014.3085

[49] Khaibullin, T., Ivanova, V., Martynova, E., Cherepnev, G., Khabirov, F., Granatov, E., Rizvanov, A., & Khaiboullina, S. (2017). Elevated Levels of Proinflammatory Cytokines in Cerebrospinal Fluid of Multiple Sclerosis Patients. Frontiers in Immunology, 8. 10.3389/fimmu.2017.00531

[50] Wedel, S., Martic, I., Guerrero Navarro, L., Ploner, C., Pierer, G., Jansen-Dürr, P., & Cavinato, M. (2023). Depletion of growth differentiation factor 15 (GDF15) leads to mitochondrial dysfunction and premature senescence in human dermal fibroblasts. Aging Cell, 22(1), e13752. 10.1111/acel.13752

[51] Vadasz, Z., Haj, T., Kessel, A., & Toubi, E. (2013). Age-related autoimmunity. BMC Medicine, 11(1). 10.1186/1741-7015-11-94

[52] He, J., Zhou, J., Yang, W., Zhou, Q., Liang, X., Pang, X., Li, J., Pan, F., & Liang, H. (2017). Dexamethasone affects cell growth/apoptosis/chemosensitivity of colon cancer via glucocorticoid receptor α/NF-κB. Oncotarget, 8(40), 67670–67683. 10.18632/oncotarget.18802

